# Evaluating the effects of aging on biodistribution and gene silencing activity of lipid-siRNA conjugates delivered into cerebrospinal fluid

**DOI:** 10.64898/2025.12.17.693452

**Authors:** Alexander P. Ligocki, Alexander G. Sorets, Adam M. Abdulrahman, Nora Francini, Joshua C. Park, Ju Ha Lee, William T. Ford, Sarah M. Lyons, Emma L. Fritsch, Zachary E. Lamantia, Plamen P. Christov, Craig L. Duvall, Ethan S. Lippmann

**Author notes:** Correspondence to: Ethan Lippmann;, Craig Duvall.

## Abstract

Aging is the primary risk factor for chronic neurodegenerative diseases and is associated with alterations to cerebrospinal fluid (CSF) flow and clearance. CSF delivery is currently the most clinically advanced route of administration for oligonucleotide therapeutics, but it remains poorly understood how aging, which is rarely incorporated into clinical trials, impacts biodistribution, gene silencing activity, and potential toxicity of these compounds. Here, we evaluated a lipid-siRNA conjugate (L2-siRNA) for potential age-related changes to CSF-mediated delivery, mRNA silencing, and safety. We found that L2-siRNA exhibited comparable biodistribution and on-target silencing of Huntingtin (*Htt*) between young and aged mice in all tested regions of the central nervous system (CNS) and across extended time points. Examining transport along CSF efflux routes revealed uptake in deep cervical lymph nodes and dura. Further, L2-siRNA did not generate detectable toxicity in the CNS or periphery of aged mice. A subset of studies benchmarked L2-siRNA against a C16 lipid-siRNA conjugate that recently entered clinical trials. Collectively, these results provide valuable insight into siRNA conjugate biodistribution and activity in the CNS in the context of aging and further establish the performance of L2-siRNA under conditions relevant to the treatment of neurodegenerative diseases.

## Introduction

Aging is the strongest known risk factor for numerous neurological disorders, including sporadic Alzheimer’s and Parkinson’s disease (1). These conditions share several physiological hallmarks such as chronic low-grade neuroinflammation, increased cellular stress, and altered cerebrospinal fluid (CSF) dynamics (2–4). However, the influence of aging on drug delivery, distribution, and efficacy is not often assessed. Instead, therapeutic compounds are typically evaluated in young adult animal models, which may not recapitulate the changes in transport and gene silencing activity that could occur in aging. As such, therapeutics that are predicted to succeed in young cohorts could potentially fail when targeting age-associated neurodegenerative disorders during clinical trials (5–7). Thus, incorporating aged models into preclinical development is essential for accurately assessing the therapeutic potential of central nervous system (CNS)-targeted drugs under physiologically relevant conditions (8).

Short-interfering RNA (siRNA) therapies are one example of a biotechnology that is gaining clinical interest for treating neurodegenerative diseases. siRNA has emerged as an attractive therapeutic strategy because it can be engineered for selective silencing of disease-associated genes, including mutant variants, by promoting degradation of target mRNA and ultimately reducing pathogenic protein levels (9). Unlike small molecules and monoclonal antibodies, which rely on potentially inaccessible or non-selective protein epitopes, siRNA offers a sequence-specific approach for mediating degradation of the mRNA rather than targeting the mature protein (9, 10). Chemical modifications added to siRNA structures limit toxicity and protect against nuclease-mediated degradation, improving target gene silencing potency and longevity (9, 11, 12). Conjugation of siRNA to peptides, antibodies, or lipids can further improve biodistribution, pharmacokinetics, and tissue-specific delivery (13–15). The FDA approval of GalNAc-conjugated siRNA drugs such as Vutrisiran and Givlaari, which target the liver via systemic administration, highlights the translational success of such strategies (16). Yet, extending this success to the CNS remains a challenge, partly due to the restrictive nature of the brain barriers (17). As such, the most direct and widely used routes for CNS-targeted siRNA delivery are intrathecal or intraventricular (ICV) administration into the CSF, and several siRNA-based drugs utilizing these delivery routes are currently under clinical investigation for neurodegenerative diseases including Alzheimer’s disease. However, given that the preclinical studies for these drugs were conducted in young animals, it remains unclear if age-related physiological changes will alter distribution and efficacy of siRNAs and affect clinical outcomes.

As CSF is the primary delivery medium for siRNAs, it is important to consider the substantial age-associated changes that occur across multiple facets of CSF physiology, including its production, composition, movement, and efflux (2, 18–20). Mice and humans exhibit a substantial decrease in CSF production in normal aging (21). Paradoxically, CSF volume increases with age, potentially due to impaired glymphatic and lymphatic drainage, which may in turn lead to prolonged availability of therapeutics within the CSF (22, 23). However, delivery to deep brain regions may be impacted due to reduced flow within the perivascular spaces (PVS) (24–26). Further, alterations in drug distribution and clearance may contribute to the elevated incidence of adverse drug reactions observed in clinical trials (27). Chronic low-grade inflammation associated with aging may compound these risks by priming the immune system for exaggerated responses to delivered therapeutics (4, 27). Overall, these considerations motivate the characterization of siRNA properties in aged models, emphasizing the need for preclinical drug delivery studies to account for age-associated changes in physiology.

To improve distribution and activity of siRNAs throughout the brain, we previously engineered a lipid-siRNA conjugate (termed “L2-siRNA”) that exhibits a balance between transport and cell uptake after administration into CSF (28). In prior work, we showed that L2-siRNA could achieve on-target silencing activity in diverse brain regions out to at least 5 months following a single ICV injection, without detectable toxicity. To build on these studies that were performed in young (3-month-old) mice, in this current work, we provide direct pharmacokinetic and knockdown comparisons to aged mice (21-month-old) using the ICV delivery paradigm. In addition, we examined distribution to canonical sites of CSF efflux such as deep cervical lymph nodes and dura, which allowed us to more comprehensively characterize L2-siRNA clearance from the CNS. Finally, we benchmarked L2-siRNA against the C16 lipid-siRNA conjugate, currently in clinical trials, using two different siRNA sequences and chemistries (13).

Collectively, this work provides detailed comparison of lipid-siRNA conjugate delivery, safety, and activity across age groups, offering critical insights into potential therapeutic performance under disease-relevant physiological conditions.

## Methods

### siRNA preparation

#### Synthesis of oligonucleotides

Oligonucleotide syntheses were performed using methods previously described (28). Sequences for all synthesized oligonucleotides are located in Supplemental Table 1. Oligonucleotides were synthesized by solid-phase methods on a MerMade 12 synthesizer (BioAutomation) using 2′-F and 2′-OMe phosphoramidites (Glen Research) and standard protecting groups. A 5′-(E)-vinylphosphonate (VP) was introduced on the antisense strand with a POM-vinyl phosphonate 2′-OMe-uridine amidite (LGC Genomics). Cy5-labeled oligos were prepared on Cy5-functionalized CPG supports, while all others used universal supports. Phosphoramidites were dissolved at 0.1 M in acetonitrile, except 2′-OMe uridine, which required 20% dimethylformamide.

#### Cleavage and deprotection of oligonucleotides

Unconjugated and conjugated sense strands were cleaved and deprotected in a 1:1 mixture of ammonium hydroxide (28–30%) and 40% aqueous methylamine (AMA) for 2 h at room temperature. Cy5-labeled oligonucleotides were processed in ammonium hydroxide alone (28–30%) for 20 h at room temperature. VP-containing antisense strands were treated with 3% diethylamine in ammonium hydroxide (28–30%) for 20 h at 35 °C.

#### Purification and characterization of oligonucleotides

Following cleavage and deprotection, oligonucleotides were dried under vacuum (Savant SpeedVac SPD120, Thermo Fisher), resuspended in nuclease-free water, and purified by HPLC (Waters 1525 EF) using a Clarity Oligo-RP column (Phenomenex) under a linear gradient [60% mobile phase A: 50 mM triethylammonium acetate (TEAA) in water; to 90% mobile phase B: methanol]. Cy5-labeled and unconjugated sense strands were first desalted (Gel-Pak, Glen Research) and further purified by reverse-phase HPLC using a linear gradient (85% to 40% mobile phase A). Fractions were dried, resuspended in nuclease-free water, sterile filtered, and lyophilized. The dimethoxytrityl (DMT) protecting group was removed from unconjugated sense strands using 20% acetic acid for 1 h at room temperature, followed by desalting.

Antisense strands were purified by anion-exchange chromatography on a 10 × 150 mm Source 15Q column (Cytiva) using a sodium perchlorate gradient (buffer A: 10 mM sodium acetate in 20% acetonitrile; buffer B: 1 M sodium perchlorate in 20% acetonitrile). Runs were performed with a 90% to 70% buffer A gradient over 30 min at 5 ml/min. Purified strands were desalted, sterile filtered, and lyophilized.

Oligonucleotide identity was confirmed by LC–MS (LTQ Orbitrap XL, Thermo Fisher) using a Waters XBridge Oligonucleotide BEH C18 column and a linear gradient (85% mobile phase A: 16.3 mM triethylamine, 400 mM hexafluoroisopropanol; to 90% mobile phase B: methanol) at 45 °C for 10 min.

### C16 amidite synthesis

For studies utilizing the C16 conjugate, a uracil phosphoramidite was synthesized with the C16 structure by the Vanderbilt Molecular Design and Synthesis Center and validated by NMR (Fig. S1). The resulting phosphoramidite was incorporated into oligonucleotide syntheses as described above. The synthesis procedure is outlined below.

#### Chemistry general

All NMR spectra were recorded at room temperature on a 400 MHz AMX Bruker spectrometer. 1H chemical shifts are reported in δ values in ppm downfield with the deuterated solvent as the internal standard. Data are reported as follows: chemical shift, multiplicity (s = singlet, d = doublet, t = triplet, q = quartet, br = broad, m = multiplet), integration, coupling constant (Hz). Normal phase purification was performed with Combi-flash Rf (plus-UV) Automated Flash Chromatography System. Solvents for reactions, extraction, and washing were ACS reagent grade.

#### Synthesis of 1-(2,2-dibutyl-6-(hydroxymethyl)tetrahydrofuro[3,4-d][1,3,2]dioxastannol-4-yl)pyrimidine-2,4(1H,3H)-dione

In a dry flask was added 1-(3,4-dihydroxy-5-(hydroxymethyl)tetrahydrofuran-2-yl)pyrimidine-2,4(1H,3H)-dione (2 gr, 8.2 mmol). Anhydrous methanol (50 mL) was added followed by dibutylstannanone (2 g, 8.2 mmol, 1 eq.) and the reaction mixture was refluxed for 2 h. The reaction mixture cooled down and concentrated. The compound was used in the next step without purification. Yield 3.7 gr (95%, purity >95%). ^1^H-NMR (400 MHz, DMSO-*d*6): δ 7.80 (d, 1H, J = 8Hz), 5.61 (d, 1H, J = 8Hz), 5.64 (d, 1H, J= 8Hz), 4.92 (tr, 1H, 8Hz), 4.04 (tr, 1H, 4Hz), 3.68-3.51 (m, 3H), 1.60-1.51 (m, 4H), 1.33-1.27 (m, 4H), 1.18-1.08 (m, 4H), 0.86 (tr, 6H, J= 8Hz).

#### Synthesis of 1-(3-(hexadecyloxy)-4-hydroxy-5-(hydroxymethyl)tetrahydrofuran-2-yl)pyrimidine-2,4(1H,3H)-dione

1-(2,2-dibutyl-6-(hydroxymethyl)tetrahydrofuro[3,4-d][1,3,2]dioxastannol-4-yl)pyrimidine-2,4(1H,3H)-dione (3.7 g, 7.78 mmol) was dissolved in anhydrous DMF (40 mL) and tetra-butyl ammonium iodide (0.28 g, 0.78 mmol, 0.1 eq.) and 1-bromohexadecane (4.75 g, 15.56 mmol, 2 eq.) were sequentially added. The reaction mixture was heated at 130°C for 8 h. The reaction mixture was diluted with ethyl acetate and washed with brine. The organic layers were dried with anhydrous MgSO4, the solid was filtered off and the solution was concentrated down. The residue was purified by normal phase chromatography (0-10% dichloromethane methanol) to give a mixture of inseparable regioisomers (2.5 g, 69%). ^1^H-NMR (400 MHz, DMSO-*d*6): δ 11.31 (s, 1H), 7.92 (d, 0.6H, J = 8Hz), 7.87 (d, 0.4H, J = 8Hz), 5.82 (d, 0.6H, J = 4Hz), 5.74 (d, 0.4H, J = 8Hz), 5.65 (tr, 0.6H, J = 2Hz) 5.63 (tr, 0.4H, J = 2Hz), 5.31 (d, 0.4H, J= 8Hz), 5.12 (overlapping tr, 1H, 8Hz), 5.03 (d, 0.6 H, J = 8Hz), 4.11-4.04 (m, 1H), 3.93-3.88 (m, 0.4H), 3.87-3.83 (m, 1H), 3.75(tr, 0.4H, 4Hz), 3.68-3.51 (m, 3H), 3.49-3.38 (m, 1H), 1.56-1.42 (m, 2H), 1.36-1.14 (m, 24H), 0.85 (tr, 3H, J= 8Hz).

#### Synthesis of 1-(5-((bis(4-methoxyphenyl)(phenyl)methoxy)methyl)-3-(hexadecyloxy)-4-hydroxytetrahydrofuran-2-yl)pyrimidine-2,4(1H,3H)-dione

1-(3-(hexadecyloxy)-4-hydroxy-5-(hydroxymethyl)tetrahydrofuran-2-yl)pyrimidine-2,4(1H,3H)-dione (mixture of regiosomers) (2.5 g, 5.3 mmol) was co-evaporated with anhydrous pyridine (3x 20 mL) and dried on high vacuum. The solid was dissolved in dry pyridine (20 mL) and DIPEA (2 ml, 11.6 mmol) and DMTrCl (1.98 g 5.87 mmol) were added. The reaction mixture was stirred at RT for 2 h. The solvent was removed in vacuum, and the residue was diluted with ethyl acetate and washed with brine. The organic layers were dried with anhydrous MgSO4, the solid was filtered off and the residue was concentrated down. The residue was purified by normal phase chromatography (0-50% hexane-ethyl acetate, 0.1% TEA) to give the desired regiosiomer (upper spot, 2 g, 50%). ^1^H-NMR (400 MHz, DMSO-*d*6): δ 11.36 (s, 1H), 7.72 (d, 1H, J = 8Hz), 7.48 (d, 2H, J = 8Hz), 7.32 (tr, 1H, J = 8Hz), 7.28-7.23 (m, 5H), 6.96-6.86 (m, 5H), 5.80 (d, 1H, J = 8Hz) 5.28 (d, 1H, J = 8Hz), 5.12 (d, 1H, J= 8Hz), 4.17 (q, 1H, J1= 8Hz, J2= 12Hz), 3.99-3.93 (m, 1H), 3.90 (tr, 1H, J= 4Hz), 3.75 (s, 6H), 3.64-3.47 (m, 2H), 3.29-3.18 (m, 2H), 1.54-1.46 (m, 2H), 1.36-1.14 (m, 28H), 0.85 (tr, 3H, J= 8Hz). ^13^C-NMR (100 MHz, DMSO-*d*6): δ 163.329, 158.541, 150.679, 145.011, 140.536, 135.436, 135.457, 130.155, 128.270, 128.270, 128.095, 127.173, 113.629, 101.875, 87.390, 86.309, 83.112, 81.248, 70.185, 68.888, 63.075, 55.419, 31.671, 29.429, 29.200, 29.085, 25.784, 22.468, 14.303

#### Synthesis of 2-((bis(4-methoxyphenyl)(phenyl)methoxy)methyl)-5-(2,4-dioxo-3,4-dihydropyrimidin-1(2H)-yl)-4-(hexadecyloxy)tetrahydrofuran-3-yl (2-cyanoethyl) diisopropylphosphoramidite

To a stirred suspension of 1-(5-((bis(4-methoxyphenyl)(phenyl)methoxy)methyl)-3-(hexadecyloxy)-4-hydroxytetrahydrofuran-2-yl)pyrimidine-2,4(1H,3H)-dione (1 g, 1.29 mmol) and 1-methyl-imidazole (0.14 g, 1.67 mmol) in anhydrous CH2Cl2 (15 mL) was added DIPEA (0.9 ml, 5.1 mmol) and 2-cyanoethyl *N*,*N*,*N*′,*N*′-tetraisopropylphosphorodiamidite (0.4g, 1.67 mmol) and stirred for 2 h. The solvent was removed in vacuum, and the residue was purified by normal phase chromatography (0-50% hexane-ethyl acetate, 0.1% TEA), (1 gr, 80%). ^1^H-NMR (400 MHz, ACN-*d*3): δ 8.96 (s, 1H), 7.83 (d, 0.6H, J = 8Hz), 7.74 (d, 0.4H, J = 8Hz), 7.79-4.74 (tr, 2H, J = 8Hz), 7.40-7.23 (m, 7H), 6.94-6.86 (m, 5H), 5.88-5.87 (m, 1H, 8Hz) 5.26-5.22 (m, 1H), 4.55-4.39 (m, 1H), 4.21-4.13 (m, 1H), 4.09-4.02 (m, 1H), 3.90-3.83 (m, 1H), 3.79 (s, 6H), 3.74-3.55 (m, 6H), 3.49-3.36 (m, 2H), 2.56 (tr, 1H, J= 8Hz), 1.64-1.54 (m, 2H), 1.36-1.14 (m, 26H), 1.22-1.14 (m, 12H), (tr, 3H, J= 8Hz). P-NMR δ 149.62, 192.24; ^13^C-NMR (100 MHz, DMSO-*d*6): δ 163.314, 158.587, 150.642, 144.929, 144.841, 140.565, 140.303, 135.528, 135.333, 135.267, 130.188, 128.243, 128.243, 128.107, 127.21, 119.177, 119.061, 113.594, 102.036, 101.934, 88.083, 87.624, 86.449, 82.317, 82.317, 82.141, 80.641, 80.062, 70.462, 70.253, 62.572, 62.145, 58.925, 58.744, 58.480, 58.283, 55.414, 43.075, 42.954, 42.863, 31.667, 29.515, 29.381, 59.311, 29.203, 29.161, 29.077, 25.809,24.710, 24.642, 22.464, 20.238, 20.171, 14.296.

### Animal husbandry

Adult male C57BL/6N mice (3-month; 18-month) were obtained from The National Institutes of Health (NIH) aged rodent colony and housed in an environmentally controlled facility under a 12-hour light/dark cycle with access to food and water. All procedures were conducted in accordance with protocols approved by the Institutional Animal Care and Use Committee (IACUC) at Vanderbilt University.

### Intracerebroventricular injection and euthanasia

siRNA duplex annealing was performed in 0.9% sterile saline by heating on a thermocycler to 95°C and gradually cooling to 4°C on the day prior to injection. Unlabeled compounds were concentrated to 1.5 mM (15 nmol) and Cy5-tagged siRNAs were concentrated to 1 mM (10 nmol) using a 3K Amicon Ultra spin filter (UFC500324). A Tecan plate reader was used to measure concentration by absorbance (260nm), and any necessary adjustments to concentration were made by adding saline. Mice were anesthetized with isoflurane and mounted on the stereotactic rig where they receive continuous isoflurane for the duration of the surgery. 5 mg/kg ketoprofen was administered subcutaneously and eye ointment was administered to prevent drying. Prior to incision, the scalp was sanitized with three alternating betadine and 70% ethanol scrubs. An incision along midline was made to open the scalp and hydrogen peroxide was applied to expose the bregma. Injection coordinates, with relation to bregma, were ± 1.0 mm medial-lateral, -2.3 mm dorsal-ventral, and -0.2 mm anterior-posterior. Two holes were drilled through the skull at these coordinates for bilateral injection. The syringe (Hamilton Model 701, blunt 30 G) was centered at bregma and brought to these coordinates and slowly lowered into the ventricle. Injections were performed at a rate of 1 µl/min, with a total of 5 µl per ventricle. The needle was left inserted in the ventricle for an additional 3 minutes after injection, followed by gradual retraction over 2 minutes. The scalp was then sutured shut, and mice were monitored for recovery. To maintain body temperature, mice were placed on a heating pad (37°C) during and after surgery. Mice received daily ketoprofen analgesia for 72 hours post operation. At the terminal timepoint, mice were euthanized by ketamine (450 mg/kg)/xylazine (50 mg/kg) overdose and transcardially perfused with cold 1x DPBS for peptide nucleic acid (PNA), RT-qPCR, Western blot, and flow cytometry outcome measure assessments; for histological readouts, mice were transcardially perfused with cold 1x DPBS and 4% paraformaldehyde (PFA).

### Histology

Following euthanasia, tissues were dissected and drop fixed for 24 hours in 4% PFA (for cryosectioning) or 10% formalin (for paraffin embedding). Prior to cryosectioning, tissues were washed three times in 1x PBS and then immersed in sucrose gradients for 24 hours (15%; 30%). The hemispheres were then embedded in OCT, sectioned to 30 μm (brains and spinal cord) or 10 μm (lymph node) on a cryostat, and stored at -80°C until staining. Antibody staining was performed as follows. First, sections were thawed to room temperature, a barrier was drawn using a hydrophobic pen, and sections were washed 3 times in 1x PBS. Tissues were blocked for 1 hour at room temperature in PBST (PBS and 0.3% triton X-100) plus 5% donkey serum (Sigma D9663). Slides were then incubated for either 2 hours at room temperature or overnight at 4°C with the primary antibody diluted in 1x PBS containing 5% donkey serum and 0.3% Triton-X 100. Primary antibodies included CD31 (1:100, BD Biosciences 550539), Lyve1 (1:200, R&D AF2125), AQP4-488 (1:200, Abcam ab284135), Glut1-PE (1:300, Abcam ab209449), F4/80 (1:100, Invitrogen 14-4801-81). Lectin-488 (1:100; Vector laboratories DL-1174) was also used. The sections were then washed three times in 1x PBS for 5 minutes, followed by a 1-hour room temperature incubation in the appropriate secondary antibody (1:1000 dilution in 1x PBS containing 5% donkey serum and 0.3% Triton-X 100). Sections were washed and incubated with DAPI (1:5000, Thermo Fisher Scientific D1306) for 5 minutes, washed again in 1x PBS, and then mounted under a coverslip with ProLong gold antifade reagent. Imaging was performed on a Leica epifluorescence microscope. For samples assessing the location of Cy5-tagged siRNA, where antibody staining was not required, slides were washed twice with 1x PBS and then mounted with ProLong gold plus DAPI.

Immunohistochemistry for GFAP was performed on paraffin embedded sections using an Epredia Autostainer 360. Sections were baked at 60°C for 1 hour, then deparaffinized and rehydrated through steps of xylene and ethanol. Xylene (two washes for 20 minutes each), and ethanol (100% and 95% three times for 3 minutes each). Antigen retrieval was then performed using the Expredia PT module in citrate buffer (pH 6) at 97°C. Slides were then moved to the auto stainer. Sections were blocked with Dako Protein Block (X0909) for 30 minutes and Flourescent block (Thermo Fisher 37565) for 20 minutes. Anti-GFAP primary antibody (1:1000, Dako Z0334) was diluted in Dako protein block and incubated for 1 hour. Slides were then washed and incubated in Alexa Fluor 647-conjugated secondary antibody (1:1000, Thermo Fisher Scientific A21245) for 1 hour. Sections were mounted with ProLong gold with DAPI (Invitrogen P36931) and imaged using the Aperio Versa 200 slide scanner. All quantification was done using ImageJ.

### Peptide nucleic acid hybridization assay

Upon euthanasia, tissue was harvested and biopsy punches taken from brain and spinal cord regions, and stored in RNAlater at -20°C. To prepare homogenates, biopsy punches were weighed and placed in 300 μl homogenization buffer (Thermo Fisher Scientific QS0518) plus proteinase-K (Thermo Fisher Scientific QS0511, 1:100). Tissue was disrupted using a Tissuelyzer 2.0 for 5 minutes at 30 Hz. Following homogenization, tissues were incubated at 65°C for 1 hour, then spun down at 15,000 × g for 15 minutes. Supernatant was collected and stored at -80°C. A standard curve was prepared at the same time as the homogenates by adding serially diluted L2-siRNA or C16-siRNA to untreated tissue. Samples were thawed and sodium dodecyl sulfate (component of homogenization buffer) was precipitated from 200 μl of homogenate with 20 μl of 3M potassium chloride. Samples were then centrifuged at 4,000 × g for 15 minutes. The supernatant was collected and centrifuged for an additional 5 minutes at the same speed to ensure complete removal of the precipitate. 150 μl of supernatant was then transferred to a screw cap tube containing 100 μl of hybridization buffer (50 mM Tris, 10% ACN, pH 8.8) and 2 μl of a 5 μM Cy3-labeled PNA probe (∼10 pmol/ 150 ul of sample, PNA bio). The sequences of the PNA probes are found in Supplemental Table 1. The antisense strand was annealed to the probe by heating to 90°C and then incubating at 50°C for 15 minutes. The samples were then run through a DNAPac PA100 anion-exchange column (Thermo Fisher Scientific) on an iSeries LC equipped with RF-20A fluorescence detector (Shimadzu). Mobile phases consisted of buffer A (50% acetonitrile and 50% 25 mM Tris–HCl, pH 8.5; 1 mM ethylenediaminetetraacetate in water) and buffer B (800 mM sodium perchlorate in buffer A), and a gradient was obtained as follows: 10% buffer B within 4 minutes, 50% buffer B for 1 minutes and 50% to 100% buffer B within 5 minutes. The final mass of siRNA was calculated using the area under the curve of Cy3 fluorescence from a standard curve. Data are reported as mass of antisense strand per mass of tissue.

### Flow Cytometry

#### Brain

Single cell suspensions were made from young (3-month) and aged (21-month) mouse brains that received 10 nmol of Cy5-conjugated L2-siRNA using the mouse adult brain dissociation kit from Miltenyi Biotec (130-107-677), with steps for myelin removal and red blood cell lysis according to the manufacturer’s instructions.

#### Dura

Single cell suspensions were made from young (3-month) and aged (21-month) mice that received a single 10 nmol injection of Cy5-conjugated L2-siRNA. Following euthanasia, dura and meninges were harvested by dissection of the dura from the skull and placed in digestion buffer (DMEM supplemented in 2% fetal bovine serum, 1 mg/mL collagenase 8 (Sigma-Aldrich, C2139) + 0.5 mg/mL DNase 1) at 37°C for 30 minutes, with gentle vortexing every 10 minutes. After 30 minutes, the dura was dissociated by triterating with a P1000 pipette 20 times and passed through a 35-um cell strainer. The cell suspension was centrifuged at 450xg at 4°C for 2 minutes, then washed two times in FACS buffer (0.5% BSA in 1x DPBS).

#### Flow cytometry staining

Cell suspensions were FcR-blocked for 10 minutes on ice (10 µl per sample, Miltenyi 130-092-575). Cells from each brain were split into separate tubes for antibody staining: ACSA2 for astrocytes (1:2,000; Miltenyi 130-123-284), and CD11b for immune cells (1:2,000; BD Biosciences 561689). Each sample was additionally stained with O1 (1:100, R&D systems FAB1327G), a marker for oligodendrocytes, to exclude these cells from analysis. For dura, cells were stained for CD11b (1:2000; Biotechne FAB7335P) and CD206 (1:100 Biolegend 141709). After washing, cells were resuspended in 1x DPBS with 0.5% BSA and DAPI (1:10,000, 5 mg/ml stock). Samples were then run on an Amnis CellStream flow cytometer. The gating scheme utilized samples stained with identical antibodies to the experimental groups but lacked Cy5 signal to identify Cy5+ cells (FMO controls from samples from an uninjected mouse). Single color compensation controls were run for each fluorophore.

### RT-qPCR

RT-qPCR methodology was utilized to determine mRNA silencing at 2 week and 3-month timepoints. Following euthanasia, brains were dissected, and biopsy punches were taken from each brain region and stored in RNAlater. Homogenates were prepared from biopsy punches in 350 µl of RLT buffer plus β-mercaptoethanol and processed with 5 mm stainless steel beads (Qiagen cat. No. 69989) for 5 minutes at 30 Hz (TissueLyser II). RNA was then extracted using an RNeasy plus mini kit (Qiagen 74134) according to the manufacturer’s instructions. RNA was eluted in 35 μl RNAse-free water, and the concentration and purity were measured by 230/260/280 nm absorbance on a Nanodrop 2000c spectrophotometer. cDNA was prepared at a standard RNA content for each brain region. Last, qPCR was performed by preparing a 10 µl reaction mixture (384 well plate used for qPCR on brain regions) or 20 µl reaction mixture (96 well plate for all other groups), composed of 2X master mix, water, Taqman probes, and cDNA sample. Taqman probes were Mm00478295_m1 (*Ppib*), Mm01213820_m1 (*Htt*), Mm01344233_g1 (*Sod1)*. Samples were run on a BioRad CFX Real Time PCR in duplicate (2 min @ 50°C, 10 min @ 95°C, then cycle 15 seconds @ 95°C and 1 min @ 60°C). All samples were analyzed according to standard ΔΔCt methodology. Each sample is normalized to *Ppib* as a housekeeping gene. Conventional RT-qPCR controls (no-template control and no reverse transcriptase control) were run on every plate and did not amplify.

### Serum chemistry

Mice receiving bilateral ICV injections of 15 nanomoles of L2-siRNA or vehicle control were euthanized 48 hours after injection. Blood was collected prior to perfusion, and serum was isolated with BD microtainer blood collection tubes (BD Biosciences 365967) via centrifugation at 1,000 x g for 10 minutes at 4°C. Samples were flash frozen and stored at -80°C. 150 µl of each sample was analyzed by Antech GLP.

### Cytokine quantification

After euthanasia, the cortex was biopsy punched, flash frozen, and stored at -80°C. To prepare homogenates, samples were thawed in cell lysis buffer (Biorad 171304006M) and homogenized three times, with 10 seconds of homogenization followed by 30 seconds on ice. The solution was then sonicated for 20 seconds and centrifuged at maximum speed for 10 minutes at 4°C. Supernatant was collected and BCA assay was performed to measure total protein as previously described. The samples were then diluted to 1 mg/ml and processed by Eve Technologies using the Mouse High Sensitive 18-Plex Discovery Assay according to the manufacturer’s protocol (Millipore Sigma).

### Cell culture and in vitro knockdown experiments

Lipofectamine-based *in vitro* studies were performed to validate gene silencing activity of different siRNA sequences conjugated to either L2 or C16. Following the manufacturer’s instructions, lipid-siRNA conjugates were mixed with RNAiMAX transfection reagent (Thermo 13778075), incubated for 5 minutes in Opti-MEM, and added to wells (12-well plate in triplicate) such that the final concentration generated was either 1 or 10 nM. For this reverse transfection, A7R5 cells were then added to each well at a density of 50,000 cells/ml in Opti-MEM. After 4 hours, the media was changed to Dulbecco’s Modified Eagle’s Medium containing 10% fetal bovine serum, and after 48 hours the cells were harvested for RT-qPCR.

## Results

### L2-siRNA exhibits comparable delivery in the young and aged brain

For the majority of our studies, we employed a blunt ended siRNA design with alternating 2’OMe and 2’F bases on both the sense and antisense strands, creating a 20-mer blunt ended ‘zipper’ design with phosphorothioate (PS) linkages on the terminal two bases. This design confers endo- and exonuclease protection, with vinyl phosphonate (VP) attached to the 5’ antisense strand for maximal silencing (11). At the 5’ end of the sense strand, we attached the lipid conjugate composed of a divalent splitter molecule, with each arm linked to a spacer of 18 ethylene glycol (3x hexaethylene glycol repeats connected by PS linkages) and 18-carbon stearyl tails (Fig. 1A). This L2-siRNA structure was originally optimized for associations with albumin in plasma to increase circulation half-life and enhance delivery at sites with leakier blood vessels (29, 30). In follow-up work, we delivered L2-siRNA into the CSF of mice using ICV injections, which yielded robust transport throughout the brain and durable silencing activity (28). These studies set the stage to explore how aging impacts biodistribution and knockdown.

**Figure 1:**
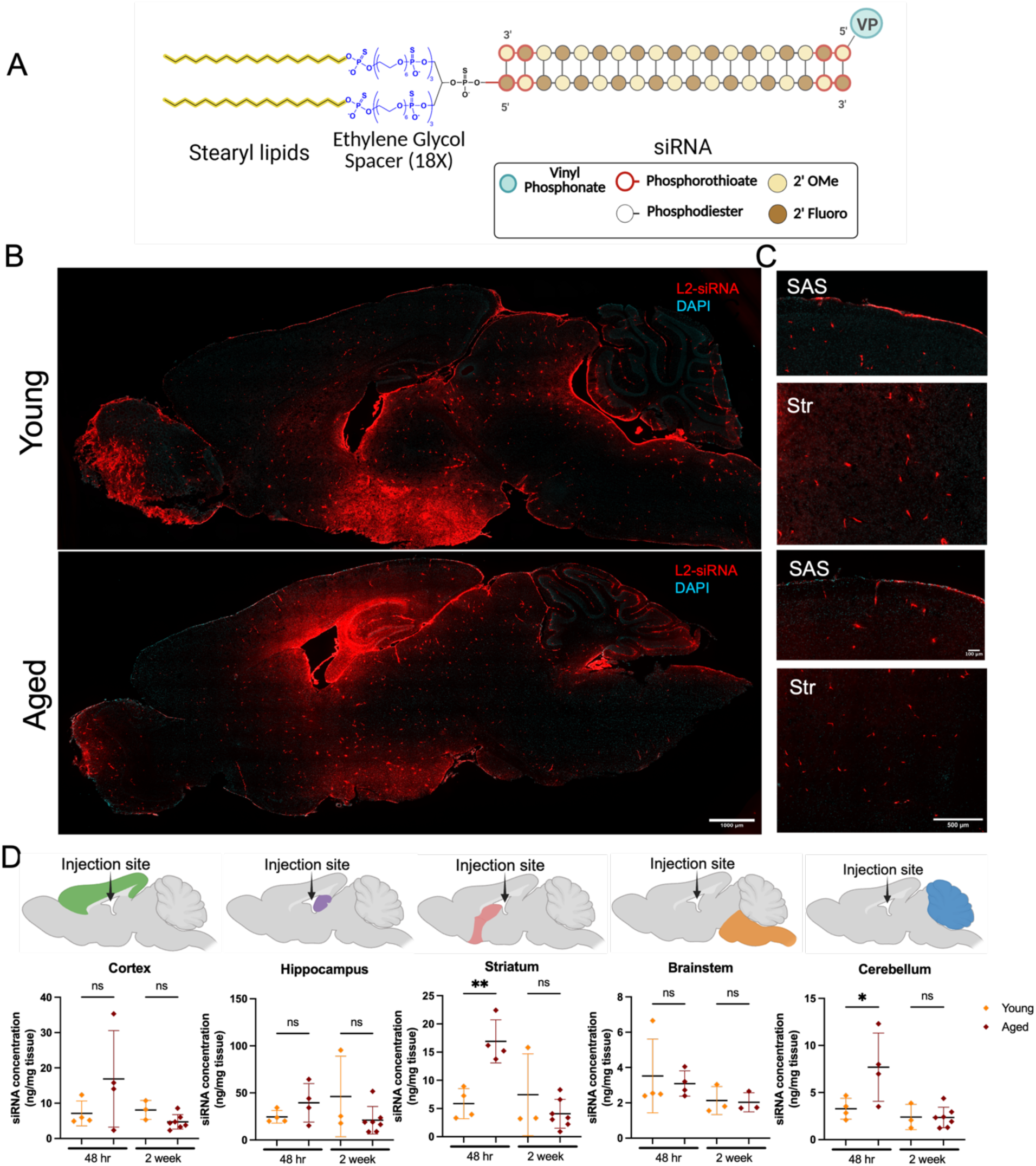
L2-siRNA biodistribution in the young versus aged brain. **A)** Diagram of L2-siRNA structure. siRNA is designed in a zipper pattern of alternating 2′ OMe, 2′Fluoro modifications with a vinyl phosphonate on the 5’ antisense strand. When used for histology, L2-siRNA also contains Cy5 fluorophore on the 3’ end of the sense strand. **B-C)** Representative images of Cy5-labeled L2-siRNA distribution in the brain, 48 hours after bilateral ICV injection of 10 nanomoles in 3-month-old (young) or 21-month-old (aged) mice. Close-ups are provided for the subarachnoid space (SAS) and striatum (Str). Patterns were consistent across N=3 mice per condition. **D)** PNA quantification of siRNA delivery 48 hours and 2 weeks after bilateral ICV injection of 15 nanomoles L2-siRNA. Each data point represents an individual mouse (N=3-7 mice per condition) and data are presented as mean ± SD. Statistical significance was determined by an unpaired t-test between young and old mice for each time point (*< 0.05, **< 0.01, ns = not significant).

We first injected young (3-month-old) and aged (21-month-old) mice using a bilateral ICV injection approach with either a total of 10 nanomoles of Cy5-labeled L2-siRNA (for histological analysis) or a total of 15 nanomoles of L2-siRNA (for PNA quantification). Tissues were collected at 48 hours (histology and PNA) or 2 weeks post-injection (PNA). To assess regional distribution, we quantified L2-siRNA in the cortex, hippocampus, striatum, brainstem, and cerebellum using a combination of microscopy and PNA-based detection. Following ICV injection, tracers have been shown to travel through the ventricular system—from the lateral ventricles to the third and fourth ventricles—before entering the subarachnoid space (SAS) via the foramina of Luschka and Magendie (Fig. 1B, C). Consistent with these established CSF flow pathways, we observed delivery of L2-siRNA to the choroid plexus (ChP) and leptomeninges, which are in direct contact with SAS CSF (Fig. 1B, C) (31). Once in the SAS, CSF penetrates the brain parenchyma via PVS surrounding penetrating blood vessels. Thus, drugs delivered into CSF will generally enter the brain parenchyma by a combination of diffusion from SAS and PVS CSF, where the latter depends on the ability of drugs to achieve convective transport through PVS flow pathways. Our prior studies found that L2-siRNA had enhanced PVS delivery compared to cholesterol-conjugated siRNA, enabling access to deep brain structures (28). Thus, we examined whether PVS-mediated distribution of L2-siRNA is preserved in the aged brain.

Histological analyses revealed similar perivascular distribution patterns in both young and aged brains. L2-siRNA signal was detected between GLUT1-positive endothelial cells and AQP4-positive astrocytic endfeet, demarcating the PVS boundary (Fig. S2). This outcome suggests that L2-siRNA retains its ability to access deep brain regions via PVS regardless of age.

Quantification of regional siRNA delivery showed the highest levels in areas proximal to the ventricles (Fig. 1D, S3). At 48 hours post-injection, aged mice exhibited higher average L2-siRNA accumulation across all examined brain regions, with statistically significant increases in the striatum and cerebellum. However, by 2 weeks post-injection, siRNA levels declined and were comparable between young and aged groups (Fig. 1D). These findings suggest that while early L2-siRNA levels are enhanced in aged brains, long-term retention is similar across age groups.

To assess cell-type-specific delivery in the aged brain, we performed flow cytometry on dissociated brain tissue from mice 48 hours after injection with 10 nanomoles of Cy5-labeled L2-siRNA. Gates were set to exclude debris, doublets, and dead cells (indicated by DAPI+ staining). ASCA2+ astrocytes and CD11b+ myeloid cells (predominantly microglia) were analyzed for percent Cy5+ and Cy5 mean fluorescence intensity (MFI, as a metric of per cell uptake) (Fig. S4). Delivery to ACSA2+ astrocytes was similar between age groups, though aged astrocytes exhibited higher MFI, indicating greater L2-siRNA uptake per cell (Fig. S4). A similar pattern was observed in CD11b+ microglia: the proportion of Cy5+ microglia was comparable between young and aged brains, but MFI was significantly higher in aged mice (Fig. S4). These results suggest age-dependent differences in cellular uptake profiles, where astrocytes and microglia in aged brains internalize more siRNA per cell.

### Spinal cord distribution of L2-siRNA is preserved with age

The CSF-filled SAS extends around the spinal cord in a manner similar to the brain, and ICV-injected compounds must travel further to reach this distal site. Histological analysis of Cy5-labeled L2-siRNA revealed similar distribution patterns in the cervical, thoracic, and lumbar spinal cord of both young and aged mice (Fig. 2A). In aged spinal cords, L2-siRNA exhibited a more pronounced gradient from the SAS, suggesting differences in parenchymal penetration or clearance. As in the brain, L2-siRNA distribution followed perivascular pathways, with signal localized around penetrating blood vessels characterized by GLUT1-positive endothelial cells and AQP4-positive astrocytic endfeet forming the glial limitans (Fig. 2B). We further used the PNA assay to quantify siRNA levels in the spinal cord at 48 hours and 2 weeks post-injection, where no significant differences were observed between young and aged mice (Fig. 2C). siRNA levels in aged mice appeared elevated at 48 hours but decreased by 2 weeks post-injection, whereas levels in the young mice were more consistent across time points, in line with the findings in the brain (Fig. 2C).

**Figure 2:**
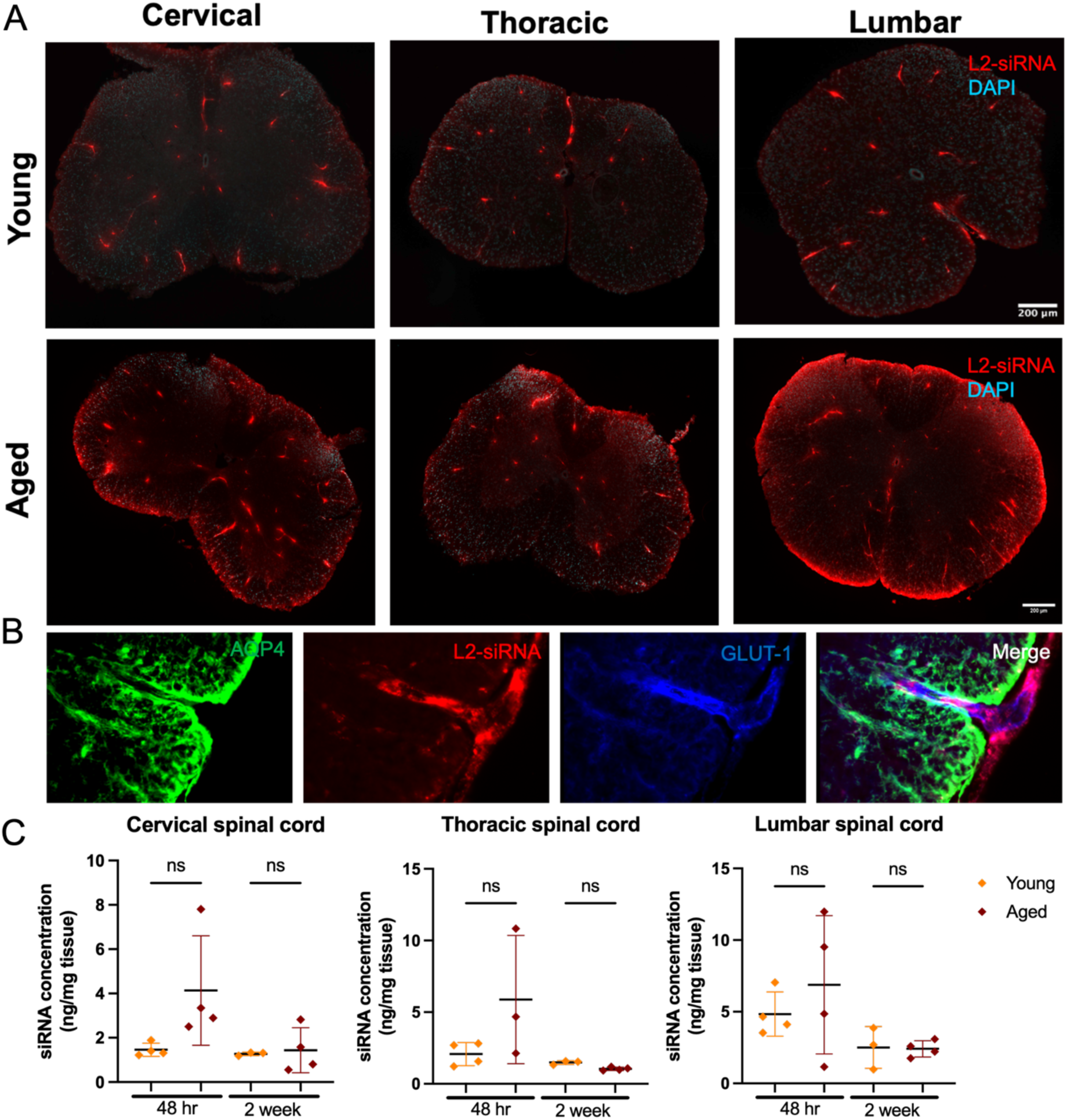
L2-siRNA biodistribution in the young versus aged spinal cord. **A)** Representative images of Cy5-labeled L2-siRNA distribution in the spinal cord, 48 hours after bilateral ICV injection of 10 nanomoles in 3-month-old (young) or 21-month-old (aged) mice. Patterns were consistent across N=3 mice per condition. **B)** Representative image of L2-siRNA distribution in the perivascular space of a penetrating blood vessel, localizing between AQP4+ astrocyte foot processes, and GLUT1+ endothelium. Patterns were consistent across N=3 mice per condition. **C)** PNA quantification of siRNA delivery 48 hours and 2 weeks after bilateral ICV injection of 15 nanomoles L2-siRNA. Each data point represents an individual mouse (N=3-4 mice per condition) and data are presented as mean ± SD. Statistical significance was determined by an unpaired t-test between young and old mice for each time point (ns = not significant).

### Aging does not diminish the gene-silencing potency of L2-siRNA targeting *Htt*

Aging is associated with chronic inflammation and impaired cellular function, both of which can reduce the efficacy of RNA-based therapeutics (32). Building on our distribution data showing minimal changes in delivery of L2-siRNA in the aged brain, we next sought to examine gene silencing activity under aged and young conditions. To assess whether L2-siRNA maintains therapeutic potency in the aged brain, we compared gene silencing activity between 3-month-old and 21-month-old mice following a single ICV injection of 15 nanomoles L2-siRNA targeting *Htt*, which encodes huntingtin and has relevance for Huntington’s disease. We used a well-validated siRNA sequence, allowing direct comparisons to prior studies (28, 33). Tissues were harvested 2 weeks post-injection to assess silencing, as earlier time points had already shown peak delivery. Knockdown of *Htt* mRNA was quantified in various CNS regions and compared to vehicle-treated controls. Robust silencing (>70%) was observed in regions proximal to the ventricles (cortex, hippocampus, and striatum) in both young and aged mice (Fig. 3A). Silencing in distal brain regions, including the cerebellum and brainstem, was also detected, though at lower levels (e.g., ∼50% in cerebellum and ∼70% in the brainstem). No statistically significant differences in knockdown were observed between young and aged mice across any brain region, with comparable average silencing across age groups. To assess the relationship between siRNA abundance and knockdown activity, we compared *Htt* mRNA silencing to siRNA delivery (quantified by PNA assay) across brain regions (Fig. 3B). Knockdown efficiency did not linearly correlate with siRNA levels, suggesting that gene silencing may reach a saturation threshold once a minimal intracellular concentration is achieved. However, based on structural and cellular heterogeneity between each region, it is possible that other factors are influencing silencing activity.

**Figure 3:**
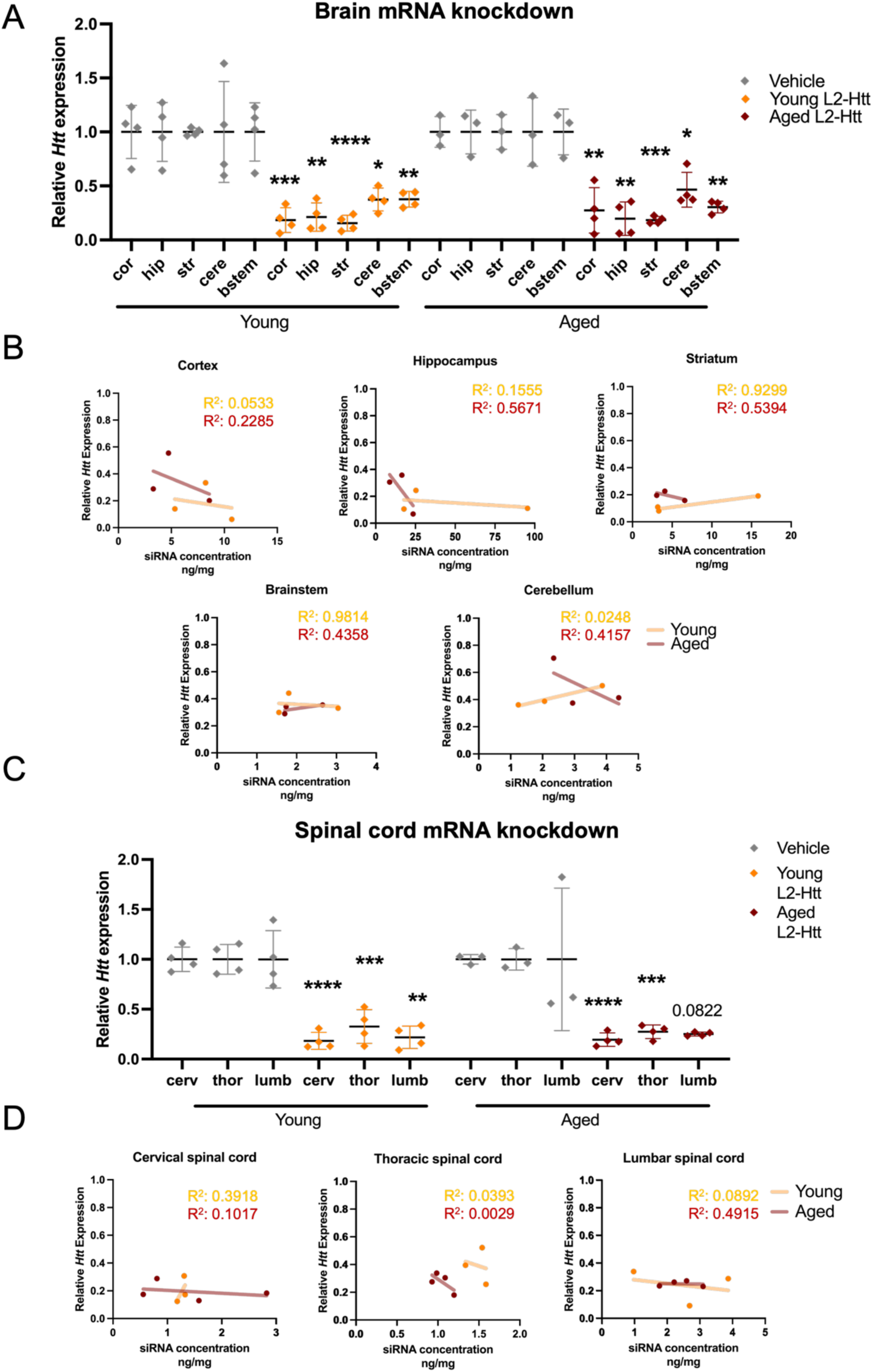
L2-siRNA exhibits robust mRNA knockdown in the aged CNS. **A)** *Htt* mRNA expression in biopsy-punched brain regions of 3-month-old (young) and 21-month-old (aged) mice, 2 weeks after bilateral ICV injection of 15 nanomoles L2-siRNA targeting *Htt* (L2-Htt) or saline. Cor = cortex, hip = hippocampus, str = striatum, cere = cerebellum, bstem = brainstem. Each data point represents a single mouse (N=3-4 mice per condition) and data are presented as mean ± SD. Statistical significance was determined by unpaired t-test for each region in each cohort between the vehicle and L2-siRNA treatment (*< 0.05, **< 0.01, ***< 0.001, ****< 0.0001, ns = not significant). **B)** Correlation between knockdown activity (data from panel A) and delivery (PNA data from Figure 1D) in brain regions. Each dot represents a biological replicate. Linearity measured by Pearsons’s correlation coefficient. **C)** *Htt* mRNA expression in cervical (cerv), thoracic (thor), and lumbar (lumb) spinal cord regions of 3-month-old (young) and 21-month-old (aged), 2 weeks after bilateral ICV injection of 15 nanomoles L2-siRNA targeting *Htt* (L2-Htt). Saline was used as a negative control. Each data point represents a single mouse (N=3-4 mice per condition) and data are presented as mean ± SD. Statistical significance was determined by an unpaired t-test (*< 0.05, **< 0.01, ***< 0.001, ****< 0.0001, ns = not significant). **D)** Correlation between knockdown activity (data from panel A) and delivery (PNA data from Figure 2C) in spinal cord regions. Each dot represents a biological replicate. Linearity measured by Pearsons’s correlation coefficient.

We next examined spinal cord regions, where significant *Htt* mRNA knockdown was detected in the cervical, thoracic, and lumbar spinal cord of young mice, and in the cervical and thoracic spinal cord of aged mice. In the aged lumbar cord, knockdown approached significance (P = 0.0659) (Fig. 3C). No statistically significant differences were observed between young and aged groups in any spinal cord region. When siRNA levels were compared to knockdown across spinal cord regions, no consistent relationship was observed (Fig. 3D), again reinforcing the idea that silencing activity is maintained above a certain delivery threshold. Overall, these findings demonstrate that L2-siRNA retains potent gene silencing activity in the aged CNS. Despite known age-related changes to CSF and parenchyma, silencing activity was comparable to young animals.

### L2-siRNA benchmarking in aged CNS

Next, we benchmarked the L2-siRNA conjugate targeting *Htt* (L2-Htt) against the C16 conjugate (C16-Htt). C16-siRNA integrates a modified phosphoramidite that places a hexadecyl chain within the interior of the siRNA structure, which improves bioavailability in the CNS after CSF delivery (13). Optimized C16-siRNAs against different targets are currently in preclinical development or phase 1 clinical trials, thus marking C16-siRNA as a useful comparator for L2-siRNA. To our knowledge, C16-siRNA has not been tested in aged animals. Thus, for these experiments, both L2-siRNA and C16-siRNA were synthesized with chemically modified blunt-ended 20-mer ‘zipper’ siRNAs as detailed earlier using the previously described *Htt*-targeting sequence. The location for the hexadecyl chain was chosen based on published design rules (13), and the potency of the C16-siRNA was confirmed *in vitro* (Fig. S5). To investigate the effect of siRNA design on potency with each conjugate, we also implemented the overhang design with enhanced stabilization chemistry (ESC) chemistry that was previously optimized for use with C16 (13). This format contains PS linkages on the terminal ends of both strands, a dinucleotide overhang on the 3’ AS and a combination of 2’F and 2’OMe ribose modifications (13).

Experiments with ESC chemistry utilized the published C16-siRNA targeting *Sod1* (C16-Sod1), and the same siRNA was synthesized in the L2-siRNA format (L2-Sod1). For knockdown studies, 18-month-old mice received an ICV injection of 15 nanomoles of L2-Htt, C16-Htt, L2-Sod1, C16-Sod1, a non-targeting L2-siRNA (L2-NTC), or saline as a vehicle control, and tissues were harvested 3 months post-injection, coinciding with 21 months of age (which matches the age utilized in prior biodistribution and knockdown measurements).

Analyses revealed that L2-Htt maintained durable mRNA silencing in all examined brain regions 3 months post-injection (Fig. 4A). C16-Htt also yielded mRNA knockdown in the cortex, striatum, and cerebellum, though to a lesser extent overall. When directly compared, L2-Htt produced significantly greater mRNA silencing in the cortex (L2: ∼80% vs C16: ∼25%) and striatum (L2: ∼50% vs C16: ∼15%) relative to C16-Htt, despite similar siRNA concentrations across regions, except in the cerebellum where C16 levels were lower (Fig. 4A, S6). Plotting relative *Htt* expression versus siRNA concentration revealed that L2-Htt retained strong silencing even in regions with low detectable siRNA, indicating durable silencing by L2-Htt (Fig. 4B). In contrast, C16-Htt showed limited silencing regardless of delivery levels. In the spinal cord, L2-Htt demonstrated robust *Htt* mRNA >50% silencing in the cervical and lumbar regions. In contrast, C16-Htt failed to induce measurable knockdown in any spinal cord region (Fig. S7).

**Figure 4:**
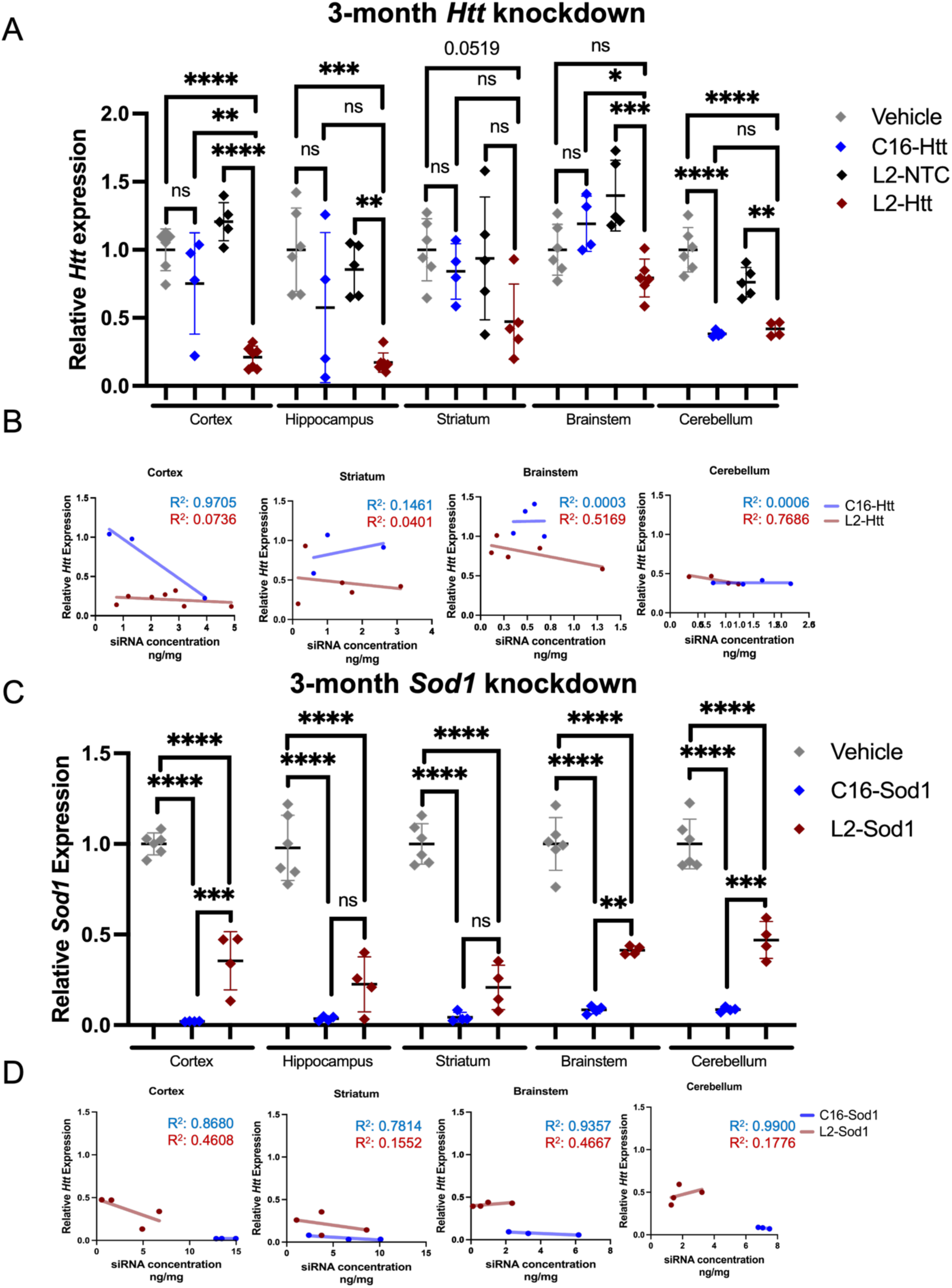
Benchmarking long-term activity of L2-siRNA in the aged CNS. **A)** *Htt* mRNA expression in biopsy-punched brain regions of 21-month-old mice, 3 months after bilateral ICV injection of 15 nanomoles L2-siRNA targeting *Htt* (L2-Htt), C16 targeting *Htt* (C16-Htt), L2-siRNA with a non-targeting sequence (L2-NTC), or saline. siRNAs were synthesized in a zipper pattern. Cor = cortex, hip = hippocampus, str = striatum, cere = cerebellum, bstem = brainstem. Each data point represents a single mouse (N=4-6 mice per condition) and data are presented as mean ± SD. Statistical significance was determined by one-way ANOVA with Tukey post hoc analysis for each brain region (*< 0.05, ***< 0.001, ****< 0.0001, ns = not significant). **B)** Correlation between knockdown activity (data from panel A) and delivery (PNA data from Supplemental Figure 5) in each brain region. Each dot represents a biological replicate. Linearity measured by Pearsons’s correlation coefficient. **C)** *Sod1* mRNA expression in biopsy-punched brain regions of 21-month-old mice, 3 months after bilateral ICV injection of 15 nanomoles L2-siRNA targeting *Sod1* (L2-Sod1), C16 targeting *Sod1* (C16-Sod1), or saline. siRNAs were synthesized with enhanced stabilization chemistry. Cor = cortex, hip = hippocampus, str = striatum, cere = cerebellum, bstem = brainstem. Each data point represents a single mouse (N=4-6 mice per condition) and data are presented as mean ± SD. Statistical significance was determined by one-way ANOVA with Tukey post hoc analysis for each brain region (**< 0.01, ***< 0.001, ****< 0.0001, ns = not significant).

L2-siRNA accumulation was also higher than C16-Htt in the thoracic and lumbar spine, whereas cervical spine delivery was not significantly different (Fig. S7).

In the context of these data, it must be stressed that the original C16 design utilized the overhang design with ESC chemistry, so it is entirely possible that the blunt-end zipper design in these experiments contributes to lower activity with the C16 conjugate. As a more direct comparison, we thus assessed silencing in aged mice using the original siRNA structure employed for the C16 conjugate, with *Sod1* as the target gene. Here, we measured potent (>90%) silencing across all brain regions with C16-Sod1, compared to 50-80% silencing with L2-Sod1 (Fig. 4C). siRNA structure additionally influenced distribution of the conjugates. Using the PNA assay, a higher concentration of C16-Sod1 was measured in the cortex, hippocampus, and cerebellum (Fig. 4D, S6). These data highlight how siRNA structure can influence conjugate delivery and knockdown. Importantly, L2-siRNA utilizing the overhang ESC chemistry still yielded significant levels of silencing, while C16 with the blunt ended zipper siRNA only elicits significant silencing in the cerebellum, suggesting the L2 lipid modification may be more potent *in vivo* with different siRNA designs. However, future comparisons to additional C16 and L2 structures will be needed to better understand these structure-function relationships.

### L2-siRNA accumulates and remains functional in the dura and deep cervical lymph nodes of aged mice

CSF is cleared through various regions that also serve as sites of immune surveillance, such as the dura and deep cervical lymph nodes (34–36). In the context of siRNA, these efflux pathways could be harnessed for therapeutic intervention or be considered for increased risk of adverse effects. How clearance of siRNAs to these sites is affected by aging is unknown, as this has not been reported in the literature previously. We measured increased levels of L2-siRNA in the aged brain at a 48 hour timepoint (Fig. 1), which can potentially be explained by delayed efflux kinetics that are known to occur in aging (37). Hence, we investigated age-related differences in the distribution and activity of L2-siRNA in the dura and deep cervical lymph nodes.

To assess biodistribution, young (3-month) and aged (21-month) mice were ICV injected with 10 nanomoles of Cy5-labeled L2-siRNA. Histology and flow cytometry were then conducted on the dura and lymph nodes after 48 hours. To assess *Htt* silencing (2 weeks) and siRNA accumulation (48 hours), mice were injected with 15 nanomoles of L2-Htt or L2-NTC. In the dura, L2-siRNA was observed colocalizing with F4/80, a marker for macrophages, primarily lining the sagittal and traverse sinuses, and to a lesser extent the non-sinus regions in young and aged mice (Fig. 5A). Using flow cytometry to quantify delivery between young and old mice, we found no significant difference in the percent Cy5+ or MFI in CD11b+ myeloid cells (Fig. S8). We did measure a significant decrease in the percentage of Cy5+ dural border-associated macrophages (CD11b+/CD206+ cells; P=0.0538), but this difference reflected a very subtle change, and no significant difference in MFI was measured (Fig. S8). When assessing bulk knockdown in the dura of aged mice, we observed ∼40% knockdown at 2 weeks compared to the non-targeting control (Fig. 5B). Because dural dysfunction has been implicated in neurodegenerative disease, our demonstration that siRNA can be delivered to and silence a target gene in the dura highlights a promising new avenue for drug delivery.

**Figure 5:**
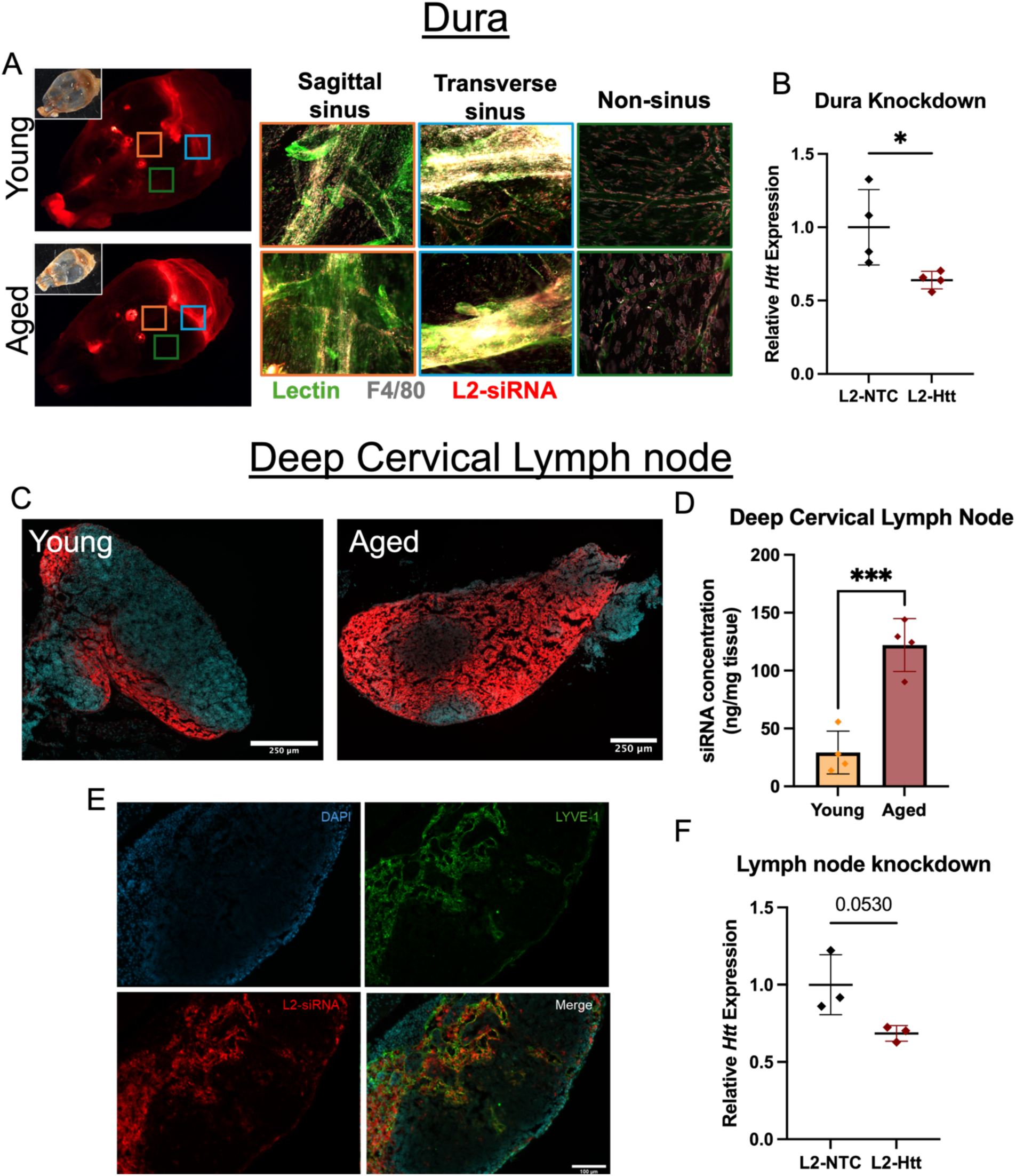
L2-siRNA accumulates and exhibits functional activity in the dura and deep cervical lymph nodes of aged mice. **A)** Representative whole mount imaging of dural distribution at the sagittal sinus, transverse sinus and non-sinus regions, 48 hours after bilateral ICV injection of 10 nanomoles Cy5-labeled L2-siRNA into 3-month-old (young) or 21-month-old (aged) mice. Blood vessels were labeled with lectin, and macrophages were labeled with the anti-F4/80 antibody. **B)** *Htt* mRNA expression in whole dura isolated from 21-month-old mice, 2 weeks after bilateral ICV injection of L2-siRNA targeting *Htt* (L2-Htt) or L2-siRNA with a non-targeting sequence (L2-NTC). Each data point represents a single mouse (N=4 mice per condition) and data are presented as mean ± SD. Statistical significance was determined by an unpaired t-test (*< 0.05). C) Representative histological images of deep cervical lymph nodes, 48 hours after bilateral ICV injection of 10 nanomoles Cy5-labeled L2-siRNA into 3-month-old (young) or 21-month-old (aged) mice. **D)** PNA quantification of siRNA delivery 48 hours after bilateral ICV injection of 15 nanomoles L2-siRNA into 3-month-old (young) or 21-month-old (aged) mice. Each data point represents an individual mouse (N=4 mice per condition) and data are presented as mean ± SD. Statistical significance was determined by an unpaired t-test (***< 0.001). **E)** Representative images of L2-siRNA co-localized with Lyve-1+ lymphatic vasculature in deep cervical lymph nodes in 21-month-old mice, 48 hours after bilateral ICV injection of 10 nanomoles Cy5-labeled L2-siRNA. **F)** *Htt* mRNA expression in deep cervical lymph nodes isolated from 21-month-old mice, 2 weeks after bilateral ICV injection of L2-siRNA targeting *Htt* (L2-Htt) or L2-siRNA with a non-targeting sequence (L2-NTC). Each data point represents a single mouse (N=3 mice per condition) and data are presented as mean ± SD. Statistical significance was determined by an unpaired t-test.

Histological examination of deep cervical lymph nodes revealed a higher degree of L2-siRNA accumulation in aging (Fig. 5C). This accumulation was primarily seen among Lyve-1+ lymphatic vessels in the aged lymph node, while young lymph node distribution was localized in the subcapsular region (Fig. 5C, D). The absolute increase in L2-siRNA accumulation was validated by the PNA assay, suggesting delayed clearance from the lymph nodes in aging (Fig. 5E) When assessing bulk knockdown in the deep cervical lymph node, we find significant (P=0.053) silencing at 2 weeks post injection compared to the non-targeting control (Fig. 5F). Collectively, these data reveal age-related L2-siRNA distribution patterns at the sites of CSF efflux, with an increased concentration in the aged deep cervical lymph nodes, while knockdown data in aged mice indicates the potential for therapeutically targeting these structures.

### Safety assessments reveal L2-Htt is well tolerated in aged mice

We previously demonstrated that L2-siRNA has a favorable safety profile at a 15 nanomole dose in young mice following ICV injection (28). These studies showed no significant changes in brain cytokine and chemokine levels, markers of gliosis, or serum chemistry indicators of systemic toxicity. To evaluate the safety of L2-siRNA in aged mice, we carried out similar experiments. Following CSF drainage, solutes enter the blood and lymphatic systems and are ultimately cleared by the liver and kidneys (38). Since age-related declines in renal and hepatic function can impair drug clearance and increase risk of toxicity (7), we began by analyzing L2-siRNA accumulation in peripheral organs. In both young and aged mice, the highest levels of L2-siRNA were detected in the liver and kidney at early time points, with subsequent decline over time, to less than 1 ng/mg by 3-months (Fig. 6A). The liver showed higher overall siRNA accumulation relative to the kidney. Aged mice further exhibited an increase in siRNA levels compared to young mice at 2 weeks post-injection, consistent with the increase observed in the lymph nodes, before declining to near baseline by 3-months. Notably, this transient hepatic accumulation was associated with ∼50% silencing of *Htt* mRNA at 2 weeks, but no significant knockdown was observed at 3 months (Fig. 6B). To further evaluate acute systemic toxicity, serum was collected 48 hours post-injection and analyzed for markers of liver (ALT, AST) and kidney (BUN, creatinine) function. No significant elevations were observed compared to vehicle-treated controls (Fig. 6C; additional markers shown in Fig. S9). As an additional safety metric, body weight was monitored for 3 months post-injection. A modest, transient weight loss was observed immediately following ICV administration of lipid-siRNA conjugates, but animals regained and maintained baseline weight over the remainder of the study (Fig. S10).

**Figure 6:**
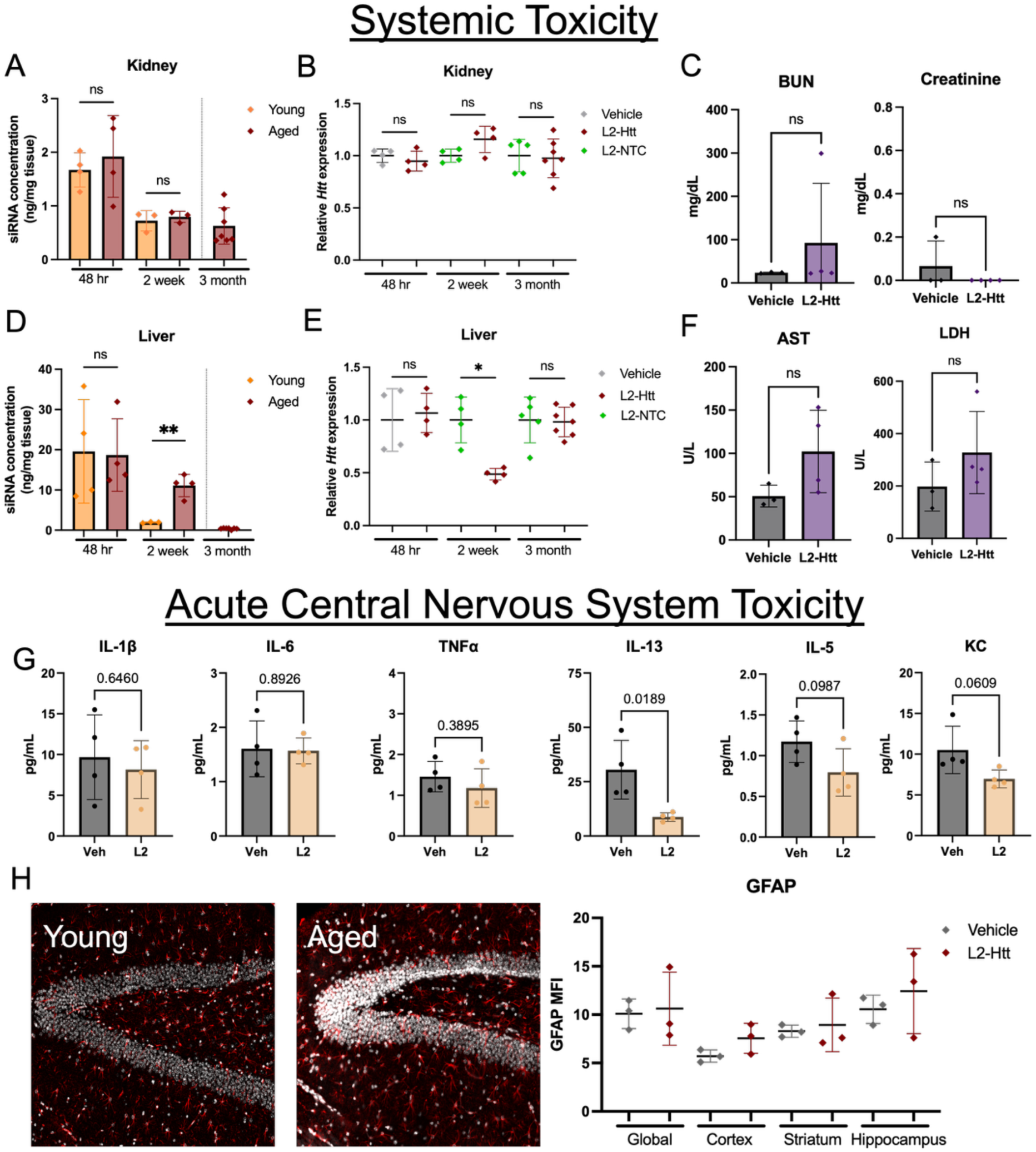
L2-siRNA does not induce toxicological signatures in aged mice. **A)** PNA quantification of siRNA levels in the kidney at defined time points following bilateral ICV injection of 15 nanomoles L2-siRNA in 3-month-old (young) or 21-month-old (aged) mice. Each data point represents a single mouse (N=3-7 mice per condition) and data are presented as mean ± SD. Statistical significance was determined by an unpaired t-test at each time point (ns = not significant). **B)** *Htt* mRNA expression in biopsy-punched kidney tissue from 21-month-old mice at defined time points following bilateral ICV injection of 15 nanomoles L2-siRNA targeting *Htt* (L2-Htt), L2-siRNA with a non-targeting sequence (L2-NTC), or saline. Each data point represents a single mouse (N=4-7 mice per condition), and data are presented as mean ± SD. Statistical significance was determined by an unpaired t-test at each time point (ns = not significant). **C)** BUN and creatinine levels in serum isolated from 21-month-old mice, 48 hours after bilateral ICV injection of 15 nanomoles L2-siRNA targeting *Htt* (L2-Htt) or saline. Each data point represents a single mouse (N=3 mice per condition) and data are presented as mean ± SD. Statistical significance was determined by an unpaired t-test (ns = not significant). **D)** PNA quantification of siRNA levels in the liver at defined time points following bilateral ICV injection of 15 nanomoles L2-siRNA in 3-month-old (young) or 21-month-old (aged) mice. Each data point represents a single mouse (N=3-7 mice per condition), and data are presented as mean ± SD. Statistical significance was determined by an unpaired t-test at each time point (**<0.01, ns = not significant). **E)** *Htt* mRNA expression in biopsy-punched kidney tissue from 21-month-old mice at defined time points following bilateral ICV injection of 15 nanomoles L2-siRNA targeting *Htt* (L2-Htt), L2-siRNA with a non-targeting sequence (L2-NTC), or saline. Each data point represents a single mouse (N=4-7 mice per condition), and data are presented as mean ± SD. Statistical significance was determined by an unpaired t-test at each time point (*<0.05, ns = not significant). **F)** AST and LDH levels in serum isolated from 21-month-old mice, 48 hours after bilateral ICV injection of 15 nanomoles L2-siRNA targeting *Htt* (L2-Htt) or saline. Each data point represents a single mouse (N=3 mice per condition) and data are presented as mean ± SD. Statistical significance was determined by an unpaired t-test (ns = not significant). **G)** Cytokine levels in biopsy-punched cortical tissue from 21-month-old mice, 48 hours following bilateral ICV injection of 15 nanomoles L2-siRNA targeting *Htt* (L2-Htt) or saline. Each data point represents a single mouse (N=4 mice per condition) and data are presented as mean ± SD. Statistical significance was determined by an unpaired t-test. **H)** Representative images of GFAP in the hippocampus and histological analysis of astrogliosis measured by mean fluorescence intensity (MFI) across the brain. Each data point represents a biological replicate (N=3 for each condition).

Aging is associated with elevated baseline levels of pro-inflammatory cytokines and reduced anti-inflammatory responses, sensitizing the brain to inflammatory stimuli (4). To assess whether L2-siRNA exacerbates neuroinflammation in aged mice, we performed cytokine/chemokine multiplex analyses on cortical samples collected 48 hours after injection. Canonical pro-inflammatory cytokines known to be elevated with age (e.g., TNF-α, IL-6, IL-1β) showed no significant increase following L2-siRNA injection relative to vehicle (Fig. 6D). Interestingly, levels of IL-13, IL-5, and KC were decreased following L2-siRNA treatment (Fig. 6D). These cytokines are typically associated with type 2 immune responses and may play anti-inflammatory or tissue-repair roles. Histological analysis of astrogliosis showed no significant changes in GFAP immunofluorescence intensity at the 2-week subacute time point (Fig. 6E). Collectively, these data demonstrate that L2-siRNA is well tolerated in aged mice, with no evidence of systemic toxicity, neuroinflammation, or organ dysfunction.

## Discussion

Aging is the predominant risk factor for neurodegeneration, yet the majority of therapeutic testing continues to rely on young animal models that fail to recapitulate the complex pathophysiology of the aged brain (39, 40). In this study, we evaluated the delivery, gene silencing activity, and safety profile of a CSF administered lipid-siRNA conjugate in aged mice. Our findings provide critical preclinical insight into the performance of siRNA therapeutics in aged models and highlight conserved activity of L2-siRNA in the aged CNS. These results further establish the potential of the L2-siRNA platform for future translation into models of neurodegenerative disease.

A major hurdle for the treatment of neurological diseases is delivery of therapeutics to desired CNS regions. Intravenous delivery of therapeutic compounds faces the highly restrictive BBB, and although the BBB becomes dysfunctional in aging, it still restricts entry of compounds into the brain (17, 41, 42). Recent work revealed conserved delivery of transferrin-receptor mediated transport of biologics across aging, despite reductions in aged mice (8). Direct CSF administration offers an alternative approach for direct CNS targeting that is especially beneficial for oligonucleotides with longer lived activity relative to proteins, but numerous age- and disease-related physiological changes to CSF composition and flow patterns may alter the pharmacokinetics and activity of candidate therapeutics (2, 18–20). L2-siRNA was originally designed to extend circulation half-life due to its reversible association with albumin, and we have also shown in prior work that L2-siRNA binds albumin in human CSF samples (28, 29). Albumin is the most abundant protein in CSF, and increased concentrations of albumin in CSF are generally measured in aging and various neurodegenerative diseases (20, 43–45). Although we have not investigated the influence of albumin binding on the distribution of L2-siRNA due to the challenges associated with such experiments, it is possible that associations with albumin contribute to a lack of significant differences in L2-siRNA activity between young and aged mice. Increased retention of L2-siRNA in the CSF due to albumin binding potentially negates decreased CSF flow and therefore lower transport throughout the CNS. We further note that C16-siRNA does not have documented albumin binding, and we did not explicitly compare C16-siRNA biodistribution and activity between young and old mice. If C16-siRNA does not bind albumin (which is likely based on its single lipid structure), it would be interesting to perform these comparisons across aging and in larger species to provide insights into whether albumin binding is an important variable in lipid-siRNA conjugate design.

CSF tracer studies have indicated a decrease in CSF-interstitial fluid exchange in aging, potentially limiting the delivery of therapeutics from CSF to aged parenchyma (46, 47). Our flow cytometry and PNA experiments indicate that L2-siRNA is effectively reaching the parenchyma and being taken up by astrocytes and microglia at comparable levels between young and aged mice, although regional differences would be masked in this bulk assay. Interestingly, using the PNA assay, we measured increased accumulation of L2-siRNA in the striatum and cerebellum of aged mice at 48 hours, which returned to similar levels as young mice by 2 weeks. However, this was not accompanied by an increase in gene silencing activity, potentially due to saturation of knockdown. We conclude that L2-siRNA can effectively reach the parenchyma and, in some regions, may even have favorable retention and activity in the aged CNS, likely due to slower fluid exchange.

CSF tracer studies have also identified age and disease-dependent deficiencies in CSF clearance through various drainage sites. For example, aged mice have delayed clearance kinetics, with reduced signal intensity of injected tracers and prolonged retention in draining cervical lymph nodes (37). Similarly, drainage to the dura mater along bridging veins and lymphatic function is compromised in aging and across diseases such as Alzheimer’s and Parkinson’s, contributing to the accumulation of neurotoxic proteins and exacerbation of neuronal damage (22, 37, 47). Our characterization of L2-siRNA clearance to these sites is therefore an important benchmark for the field. Further, neuroinflammation is generally increased in aging and disease, but we did not observe any changes to markers of systemic toxicity, cytokine levels in the cortex, or astrogliosis in aged mice receiving L2-siRNA. Thus, we infer that L2-siRNA is not causing overt toxicity in the aged CNS and is cleared through endogenous mechanisms.

Lipid-siRNA conjugates are two-component systems in which both the lipid and the siRNA are chemically modified to promote structural stability, efficient intracellular delivery, and potent gene silencing activity. However, it remains unclear whether lipids optimized for a specific siRNA structure will retain their properties when conjugated to siRNAs containing different sequences or chemical backbones. As such, we compared two platforms, L2-siRNA and C16-siRNA, by pairing each lipid with siRNAs containing either zipper or ESC modifications. This experimental design was motivated by the fact that L2-siRNA was originally developed using blunt-ended zipper design (alternating 2′-O-Me and 2′-F modifications), whereas C16-siRNA was optimized for a different ribose pattern and includes a 2-base pair overhang (13, 29). Using the zipper siRNA format, we found that L2-siRNA achieved greater *Htt* knockdown after ICV injection than C16-siRNA in several brain regions, and in some areas, C16-siRNA exhibited no significant knockdown. Notably, the internal position of the C16 lipid was not systematically varied (“walked”) to identify the optimal attachment site, although activity of the chosen C16-siRNA design was confirmed *in vitro*. In contrast to the *Htt*-zipper results, the opposite trend was observed for the Sod1-targeting ESC siRNA; C16-siRNA produced near-complete target silencing, whereas L2-siRNA activity was attenuated but still significant for each assayed region relative to the vehicle control. Overall, our findings indicate that each lipid performs best with the siRNA structure in which it was initially optimized, suggesting a sequence- or chemistry-dependent interplay between the lipid and siRNA components that warrants further investigation.

Overall, this study compares the delivery, activity, and safety of a CSF delivered lipid-siRNA conjugate in aged mice. Collectively, the results underscore the potential of L2-siRNA for the treatment of neurodegenerative diseases. Future work will focus on analyzing the efficacy of L2-siRNA in various disease models where aging is a relevant consideration.

## Acknowledgements

Figure 1A and 1D were created using BioRender and cited in the figure legend. Mass spectrometry characterization of oligonucleotides was performed in the Mass Spectrometry Research Center at Vanderbilt University. Synthesis of the hexadecyl-modified phosphoramidite C16 was provided by the Vanderbilt Institute of Chemical Biology, Molecular Design and Synthesis Center (MDSC), Vanderbilt University. Flow cytometry was performed in the VUMC Flow Cytometry Shared Resource, which is supported by the Vanderbilt Ingram Cancer Center that is funded in part by NIH grant P30 CA68485. Whole slide imaging was performed in the Digital Histology Shared Resource at Vanderbilt University Medical Center.

## Author Contributions

Conceptualization: A.P.L., C.L.D., and E.S.L. Methodology: A.P.L. performed majority of experiments with assistance from A.M.A, A.G.S., N.F., J.C.P., J.L., W.T.F., S.M.L, E.L.F., and Z.E.L. P.P.C. developed and executed the synthesis scheme for the C16 phosphoroamidite. Supervision: C.L.D., and E.S.L. Funding: C.L.D. and E.S.L. Writing—original draft: A.P.L., and E.S.L. Writing—review & editing: All authors reviewed and edited the manuscript.

## Funding

This work was supported by the National Institutes of Health grants R01 AG092015 (to E.S.L. and C.L.D.) and RF1 NS130334 (to E.S.L.). A.P.L. was supported by National Institutes of Health grant T32 ES007028. A.M.A. was supported by National Institutes of Health grant F30 AG094191.

## Disclosures

E.S.L., C.L.D., and A.G.S. are listed as inventors on patent application number 18/737720 (pending; applicant: Vanderbilt University), which relates to lipophilic siRNA conjugates for the treatment of CNS disorders.

## Data and materials availability

Correspondence and requests for material should be addressed to Ethan Lippmann at.

