## Supplemental information for "Evaluating the effects of aging on biodistribution and gene silencing activity of lipid-siRNA conjugates delivered into cerebrospinal fluid"

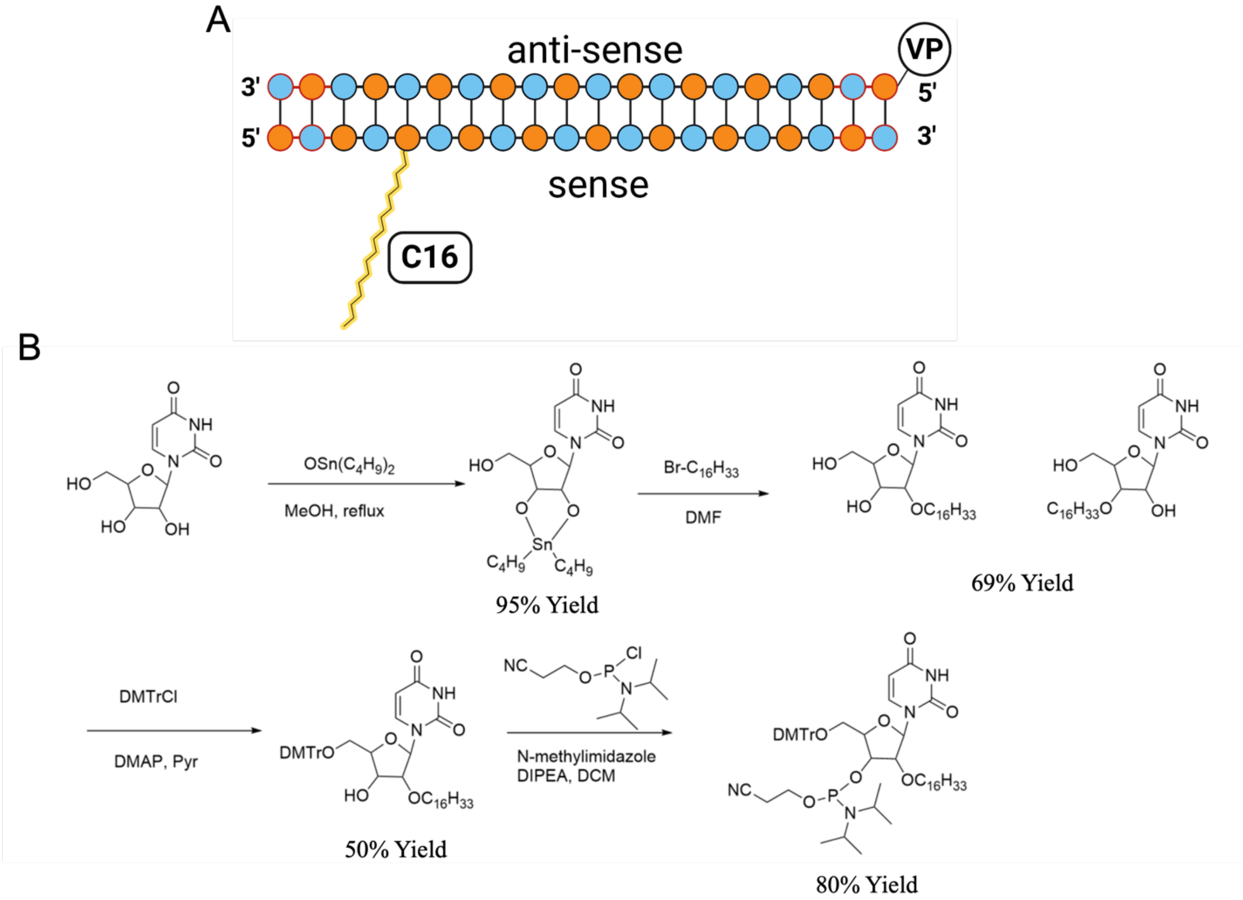


**Supplemental Figure 1: Chemical synthesis of the C16 phosphoramidite**

A) Schematic representation of C16 lipid conjugate addition onto the siRNA sense strand.

B) Representative workflow of C16 conjugate synthesis.

A

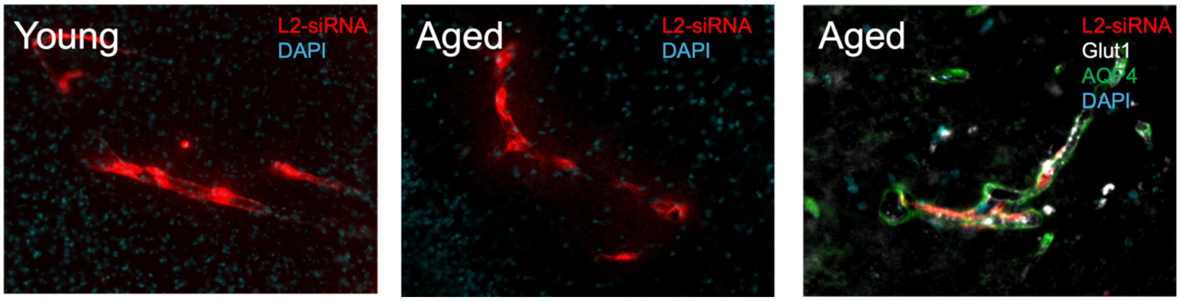


**Supplemental Figure 2: Perivascular distribution is observed in both young and aged mice.**

Representative images taken across brain regions (N=3 biological replicates) of 3-month-old (young) and 21-month-old (aged) mice, 48 hours after bilateral ICV injection of 10 nanomoles Cy5-labeled L2-siRNA. Perivascular localization of Cy5 signal was observed between astrocytic foot processes (AQP4+) and Glut1+ endothelium.

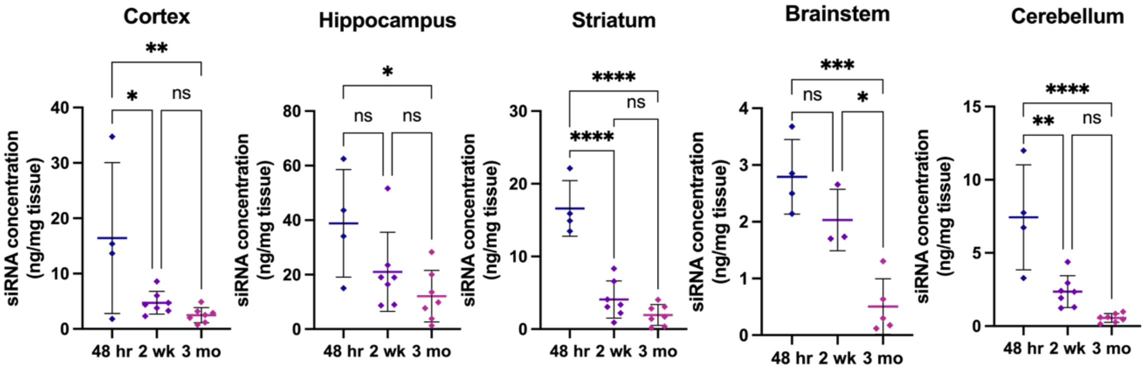


**Supplemental Figure 3: PNA analysis of L2-siRNA concentration in the aged brain over time.**

Comparison of siRNA concentrations within each region in the brain at defined time points after bilateral ICV injection of 15 nanomoles L2-siRNA$^{Htt}$. Mice received the injections at 18-months (3-month timepoint) or 21-months (48 hour and 2-week timepoint) of age. Each data point represents a single mouse (N=4-7 mice per condition) and data are presented as mean ± SD. Statistical significance was determined by one-way ANOVA with Bonferroni correction for multiple comparisons. (*< 0.05, **< 0.01, ***< 0.001, ****< 0.0001, ns = not significant).

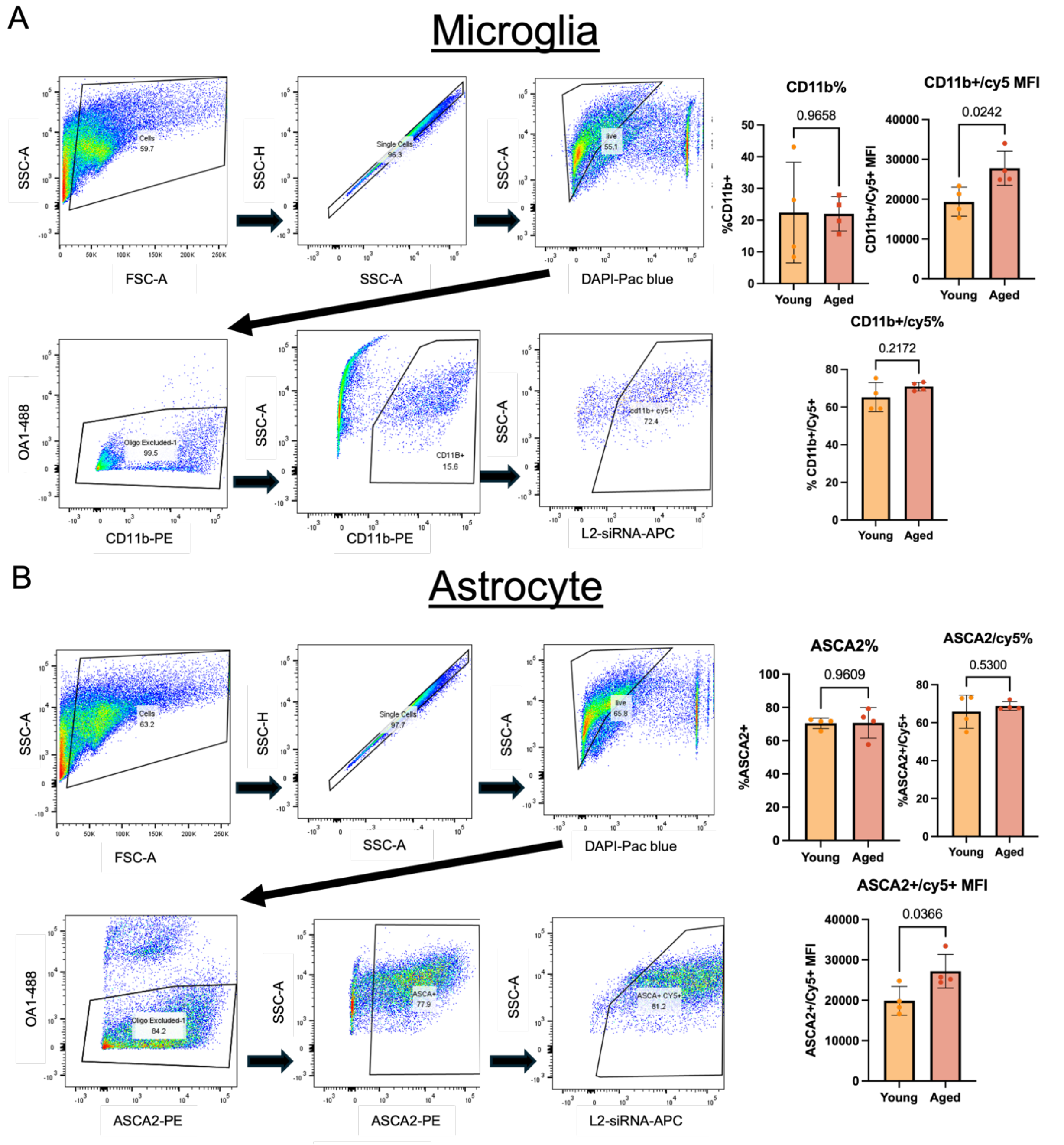


**Supplemental Figure 4: Assessment of L2-siRNA uptake in astrocytes and microglia as a function of age.**

Flow cytometry analysis 48 hours after bilateral ICV injection of 10 nanomoles Cy5-labeled L2-siRNA into young (3-month-old) and aged (21-month-old) mice. Panel A shows the gating strategy and quantifications for CD11b+ microglia, while Panel B shows the same information for ASCA2+ astrocytes. For quantification, each data point represents a single mouse (N=4 mice per condition) and data are presented as mean ± SD. Statistical significance was determined by unpaired t-test.

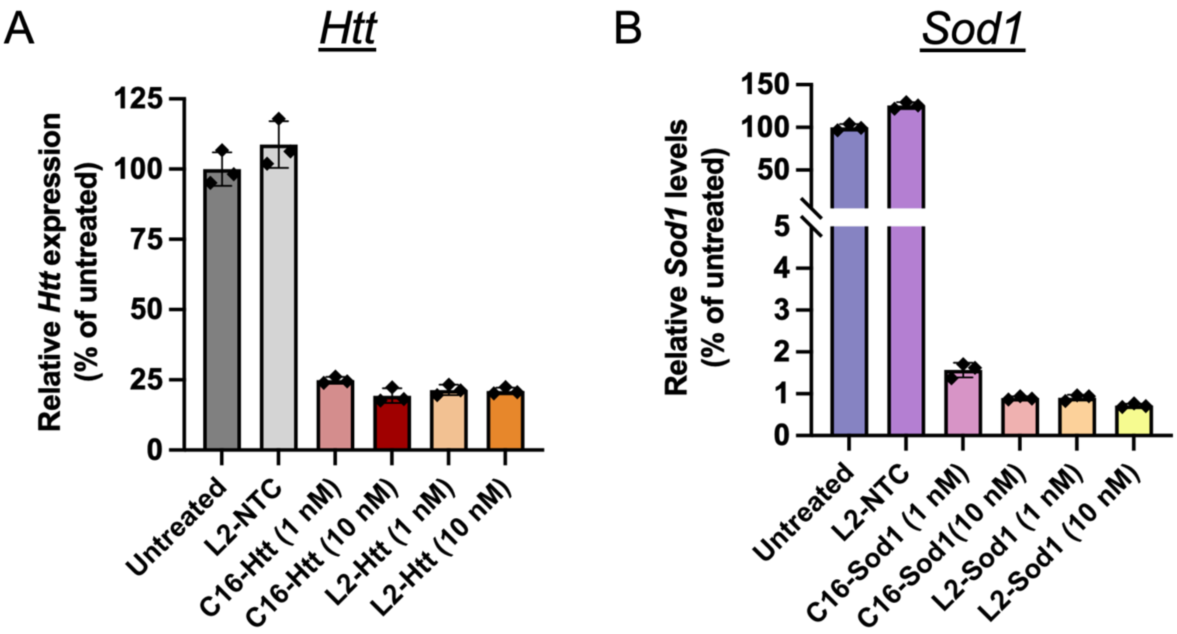


**Supplemental Figure 5:** ***In vitro*** **potency of siRNA sequences.**

A) Relative mRNA levels in A7R5 cells transfected with 1 or 10 nM C16-Htt, L2-Htt, or L2-NTC. Gene expression was quantified 48 hours after RNAiMAX transfection. Each data point represents an individual well of cells (N=3 replicates per condition), and data are presented as mean ± SD normalized to the untreated control.

B) Relative mRNA levels in A7R5 cells transfected with 1 or 10 nM C16-Sod1, L2-Sod1, or L2-NTC. Gene expression was quantified 48 hours after transfection. Each data point represents an individual well of cells (N=3 replicates per condition), and data are presented as mean ± SD normalized to the untreated control.

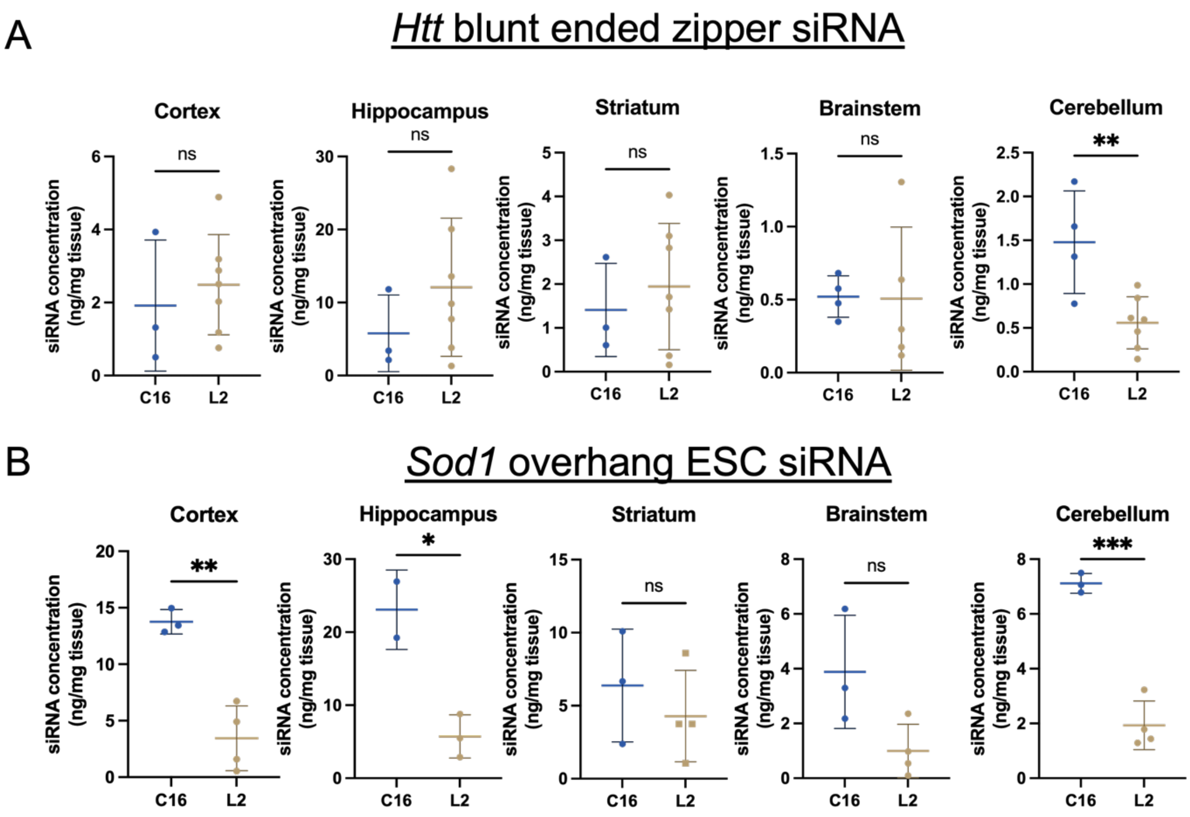


**Supplemental Figure 6: Distribution of C16-Htt, L2-Htt, C16-Sod1, and L2-Sod1 in the aged brain.**

Aged mice (18-months) received bilateral injections of 15 nanomoles of C16-Htt, L2-Htt, C16-Sod1, or L2-Sod1. *Htt*-targeting siRNAs utilized blunt-end zipper chemistry (panel A) and *Sod1*-targeting siRNAs utilized overhang ESC chemistry (panel B). siRNA levels were quantified 3 months after injection using the PNA assay. Each data point represents an individual mouse (N=3-7 mice per condition) and data are presented as mean ± SD. Statistical significance was determined by an unpaired t-test (*< 0.05, **< 0.01, ***< 0.001, ns = not significant).

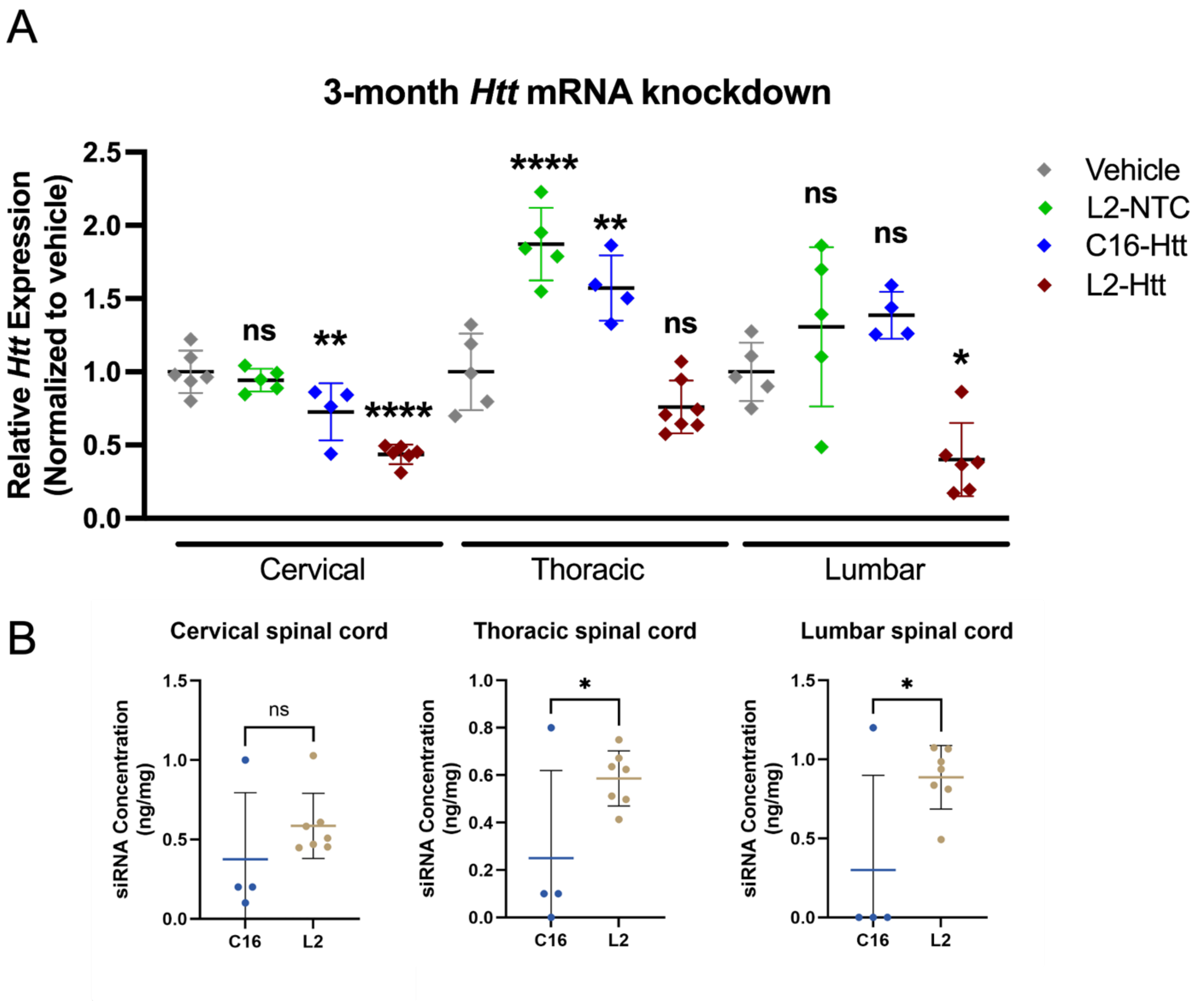


**Supplemental Figure 7: Distribution and gene silencing activity of C16-Htt and L2-Htt in the spinal cord.**

A) Aged mice (18-months) received bilateral injections of 15 nanomoles C16-Htt, L2-Htt, L2-NTC, or vehicle control. Spinal cord regions were harvested 3-months post injection and gene expression was quantified using RT-qPCR. Each data point represents an individual mouse (N=4-7 mice per condition) and data are presented as mean ± SD normalized to the vehicle control for each region. Statistical significance was determined by one-way ANOVA (*< 0.05, **< 0.01, ****< 0.0001, ns = not significant).

B) siRNA levels were quantified 3 months after injection using the PNA assay. Each data point represents an individual mouse (N=4-7 mice per condition) and data are presented as mean ± SD. Statistical significance was determined by an unpaired t-test (*< 0.05, ns = not significant).

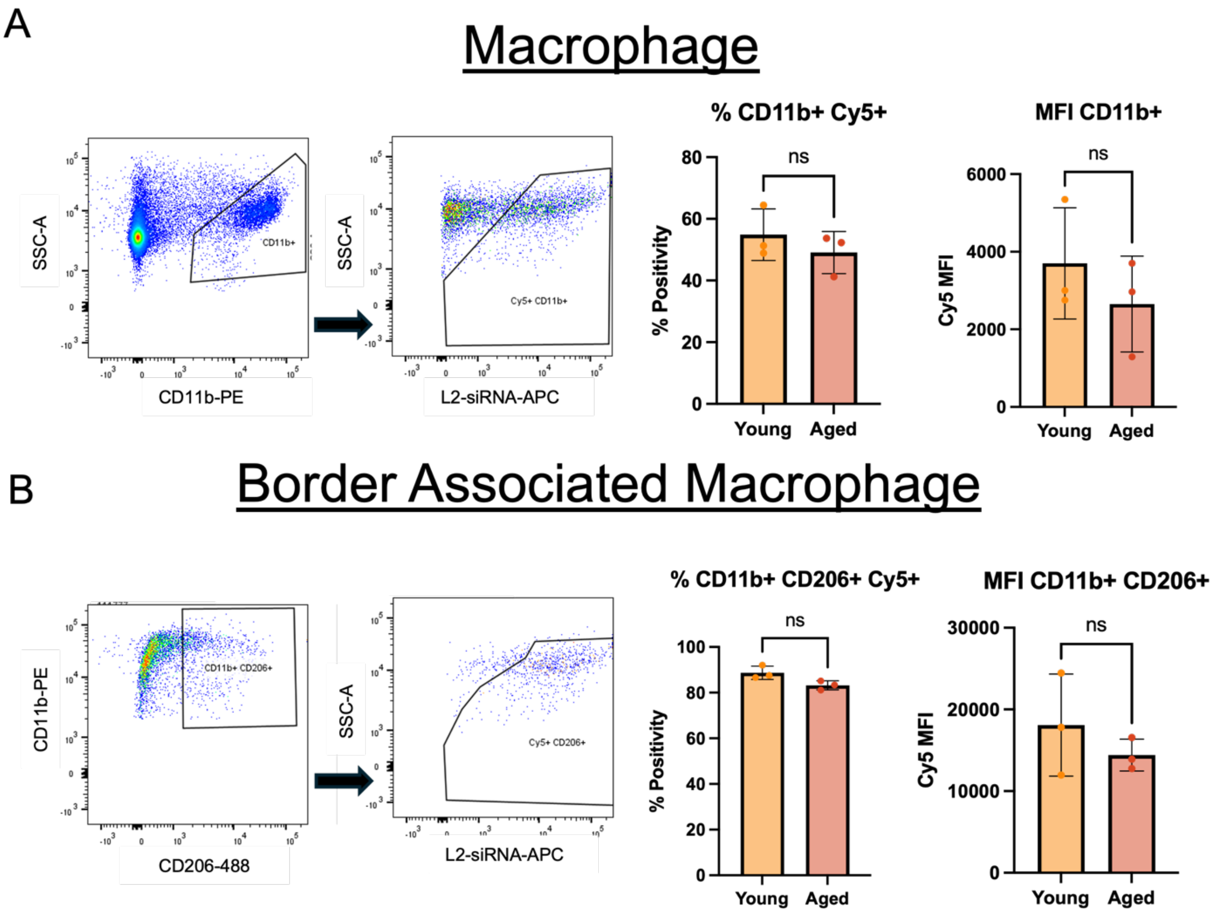


**Supplemental Figure 8: L2-siRNA uptake by myeloid cells in the young and aged dura.**

A) Flow cytometry analysis of myeloid cell uptake in the young and aged dura following bilateral ICV injection of 10 nanomoles of Cy5-labeled L2-siRNA. The gating strategy for CD11b and Cy5 is presented, and the graphs show percent CD11b+/Cy5+ and Cy5 MFI in CD11b+/Cy5+ cells. Each dot represents an individual mouse (N=3 mice per condition) and data are presented as mean ± SD. Statistical significance was determined by unpaired an t-test (ns = not significant).

B) Flow cytometry analysis of macrophage uptake in the young and aged dura following bilateral ICV injection of 10 nanomoles of Cy5-labeled L2-siRNA. The gating strategy for CD11b, CD206, and Cy5 are presented, and the graphs show percent CD11b+/CD206+/Cy5+ and Cy5 MFI in CD11b+/CD206+/Cy5+ cells. Each dot represents an individual mouse (N=3 mice per condition) and data are presented as mean ± SD. Statistical significance was determined by unpaired t-test (ns = not significant).

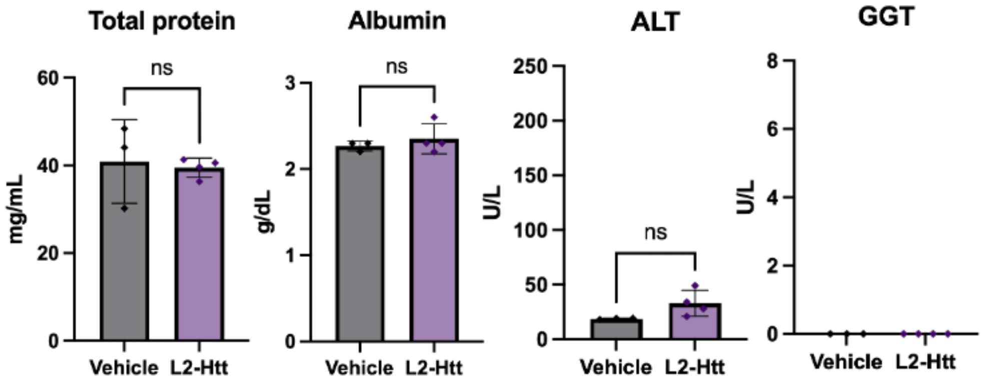


**Supplemental Figure 9: Serum analyses following ICV delivery of L2-siRNA in aged mice**.

Blood serum was isolated from 21-month-old mice 48 hours after bilateral ICV injection of 15 nanomoles of L2-Htt or saline control. Serum was sent for analysis by Antech GLP. Total protein was measured by BCA and albumin, alanine aminotransferase (ALT), and gamma-glutamyl transferase (GGT) were measured by a serum chemistry panel. Each dot represents an individual mouse (N=3-4 mice per condition) and data are presented as mean ± SD. Statistical significance was determined by an unpaired t-test (ns = not significant).

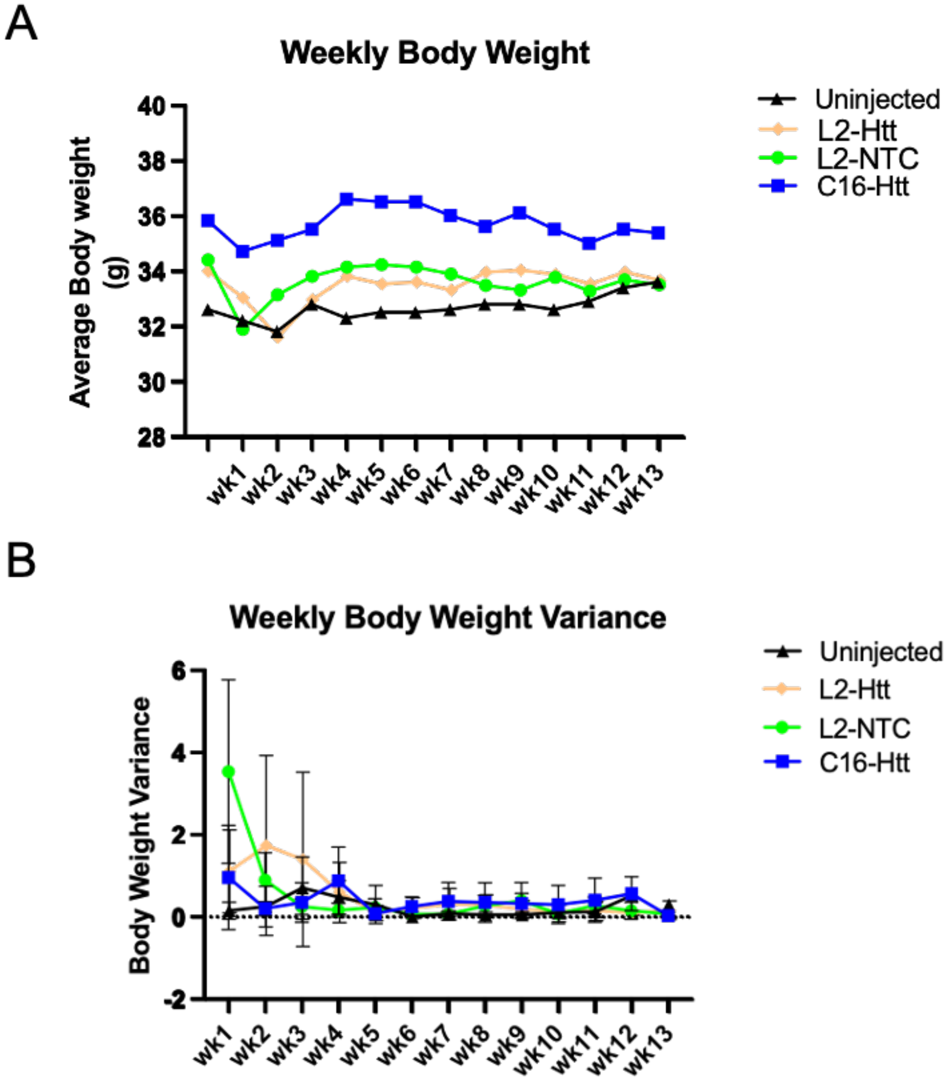


**Supplemental Figure 10: Body weights of aged mice following ICV injection.**

A) Body weights of 18-month-old mice were recorded weekly for 3-months following bilateral ICV injection of 15 nanomoles L2-Htt, L2-NTC, C16-Htt, or saline control. Each data point represents the average weight of all mice within each treatment condition (N= 3-7 per condition).

B) Variance in body weight from the prior week was recorded for 3 months post injection. Error bars represent SD within each treatment condition (N=3-7 per condition).

| Target | Sense/antisense (5’-3’) | Sequence |
|---|---|---|
| Htt | S | (MeU)*(fA)*(MeU)(fA)(MeU)(fC)(MeA)(fG)(MeA)(fA)(MeA)(fA)(MeG)(fA)(MeG)(fA)(MeU)(fU)*(MeA)*(fA) |
| | AS | VP(meU)*(fU)*(MeU)(fA)(MeU)(fC)(MeU)(fC)(MeU)(fU)(MeU)(fA)(MeC)(fU)(MeG)(fA)(MeU)(fA)*(MeU)*(fA) |
| LUC (NTC) | S | (fC)*(MeA)*(fA)(MeU)(fU)(MeG)(fC)(MeA)(fC)(MeU)(fG)(MeA)(fU)(MeA)(fA)(MeU)(fG)*(MeA)*(fA) |
| | AS | VP(MeU)*(fU)*(MeC)(fA)(MeU)(fU)(MeA)(fU)(MeA)(fU)(MeC)(fA)(MeG)(fU)(MeG)(fC)(MeA)(fA)(MeU)*(fU)*(MeG) |
| SOD1 | S | (MeC)*(MeA)*(MeU)(MeU)(MeU)(C16u)(fA)(MeA)(fU)(fC)(fC)(MeU)(MeC)(MeA)(MeC)(MeU)(MeC)(MeU)(MeA)*(MeA)*(MeA) |
| | AS | VP (MeU)*(fU)*(MeU)(MeA)(MeG)(fA)(MeG)(fU)(fG)(MeA)(MeG)(MeG)(MeA)(fU)(MeU)(fA)(MeA)(MeA)(MeA)(MeU)(MeG)*(MeA)*(MeG) |
| Htt PNA probe | | 5’N Cy3-OO-TATATCAGTAAAGAGATTAA 3’/C |
| SOD1 PNA probe | | 5'/N Cy3-OO CTC ATTTT AATCC TCACT CTAAA 3'/C |
| Legend: S = sense; AS = antisense; Me = O-methyl; * = phosphorothioate; NTC = non-targeting control; PNA = peptide nucleic acid | | |

**Supplemental Table 1:** Sequences of the sense and antisense strands of *Htt, LUC* (NTC), and *SOD1* siRNA, as well as sequences of the *Htt* and *Sod1* PNA probes.
